# Keeping SCORE enables interpretable uncertainty-aware classification from diffusion models for genomics

**DOI:** 10.1101/2025.11.26.690838

**Authors:** Benjamin Kuznets-Speck, Jaekwon Jung, Pornchanan Pholraksa, Adrianne Zhong, Leon Schwartz, Ekta Prashnani, Suriyanarayanan Vaikuntanathan, Yogesh Goyal

**Affiliations:** Department of Cell & Developmental Biology, Feinberg School of Medicine, Northwestern University, Chicago, IL, USA; Center for Synthetic Biology, McCormick School of Engineering, Northwestern University, Chicago, IL, USA; Robert H. Lurie Comprehensive Cancer Center, Feinberg School of Medicine, Northwestern University, Chicago, IL, USA; NSF-Simons National Institute for Theory and Mathematics in Biology, Chicago, IL, USA; NVIDIA, Santa Clara, CA, USA; Chan-Zuckerberg Biohub Chicago, LLC, Chicago, IL, USA; Department of Chemistry, University of Chicago, Chicago, IL, USA; Data Science of Systems Biology, Technical University of Munich, Germany; Department of Chemical & Biological Engineering, McCormick School of Engineering, Northwestern University, Evanston, IL, USA

**Keywords:** Diffusion models, Artificial intelligence, Maximum likelihood, Cell state, Genomics

## Abstract

Classifying cellular states from high-dimensional molecular and genomic measurements requires methods that provide not only accurate predictions but also calibrated uncertainty and interpretability. Current nonlinear classifiers offer accuracy but often lack uncertainty quantification and mechanistic insights into the features that matter most. We introduce Keeping SCORE, a framework that transforms conditional diffusion models into probabilistic engines for classification and regression by computing exact likelihoods along stochastic noising trajectories. We first benchmark Keeping SCORE on image recognition tasks (handwritten digits, natural photos). We then apply Keeping SCORE to single-cell transcriptomics across a 22-million-cell atlas, classifying 164 cell types with accuracy matching or exceeding state-of-the-art methods, while uniquely providing posterior probability estimates and prediction confidence. For genetic perturbation mapping across 100 CRISPRi conditions in a multi-study Perturb-seq dataset, our approach again matches or surpasses discriminative baselines, with feature-level attributions identifying which genomic features drive each decision. Applied to large-scale protein sequence data, our framework accurately regresses mutational stability effects, attributing them quantitatively to positions along the input sequence. Keeping SCORE requires no retraining or architectural changes to existing diffusion models, providing portable, interpretable, and uncertainty-aware predictions for biological discovery.

## INTRODUCTION

Classification and regression form the foundation of modern machine learning and computational biology, enabling models to sort, predict, and generalize across complex datasets [1–3]. When data are grouped into categories, a core challenge is deciding which category a previously unseen sample belongs to. Regression problems, on the other hand, like that of predicting folded protein structure from sequence aim to forecast continuous unseen quantities from auxiliary signals. From recognizing cellular phenotypes in microscopy images [4, 5] to predicting protein stability from sequence [6], or disease state from gene expression across populations [7–10], these inference tasks underpin many essential questions in computational biology and biophysics. However, it is often the case that boundaries between classes are blurred, forcing us to look beyond these dividing surfaces and into the geometry of the distributions themselves.

Conventional approaches, like linear or deep neural network regressors and classifiers, have seen great success at mapping high dimensional input data to a low dimensional set of regression parameters or classes [11, 12]. However, typical implementations remain limited in that they reduce learning to the placement of rigid decision boundaries or fitting of point estimates, overlooking the rich geometry and uncertainty of the full data distribution [13]. Specifically, most deep classification models are not equipped for uncertainty estimation, and unlike their linear counterparts, they lack interpretability, i.e. the ability to directly link input features to classification performance, for example. In recent years, generative models—particularly diffusion models—have transformed machine learning by sampling new points from the entire data manifold [14, 15]. These models have been crucial for image and molecular generation, but their capacity for inference, past ad-hoc placement of classifiers on top of diffusion models, remains largely untapped. For instance, the question of how to estimate the probability of each class, c, given a new data point x naturally within the context of diffusion models has just begun to be broached [16, 17]. Recent work has connected path-integral formulations to diffusion model likelihoods [18–22], but existing approaches either rely on approximations, target generative modeling rather than classification, or require computing a costly score divergence term.

Here, we develop a new unified framework—Keeping SCORE—that treats diffusion models as Bayesian inference machines for classification and regression (Fig. 1). Our approach eliminates the costly score divergence term exactly, yielding the first tractable and exact class-likelihood estimator for stochastic diffusion trajectories. Pretrained or newly trained diffusion models can then be used to estimate class posteriors *p*(*c*|*x*) ∝ *p*(*x*|*c*), quantify uncertainty directly from the geometry of the learned score function, and assign interpretable contributions to each input feature or degree of freedom. This framework is intrinsically uncertainty-aware, interpretable, and portable, requiring no architectural changes or retraining. The key insight is that the conditional score—the gradient of the log-probability with respect to the data—naturally encodes class probabilities through a thermodynamic path observable. We introduce this conceptual framework and demonstrate its broad applicability across domains. First, we test Keeping SCORE on classical computer vision benchmarks, including handwritten digits (MNIST [23], Fig. 1) and natural images (CIFAR10 [24]), where it provides distribution-aware classification. We then apply the framework to single-cell genomics, successfully distinguishing cell types across a 22-million-cell atlas and classifying genetic perturbations in single cells from multi-study Perturb-seq data. Finally, we extend the method to continuous regression problems—predicting changes in protein stability (ΔΔG) upon amino acid mutation—illustrating how diffusion-based inference captures quantitative biophysical effects. Together, these results establish diffusion models as general-purpose, geometry aware, inference engines that unify generative modeling, uncertainty quantification, and interpretability in genomics.

**FIG. 1.**
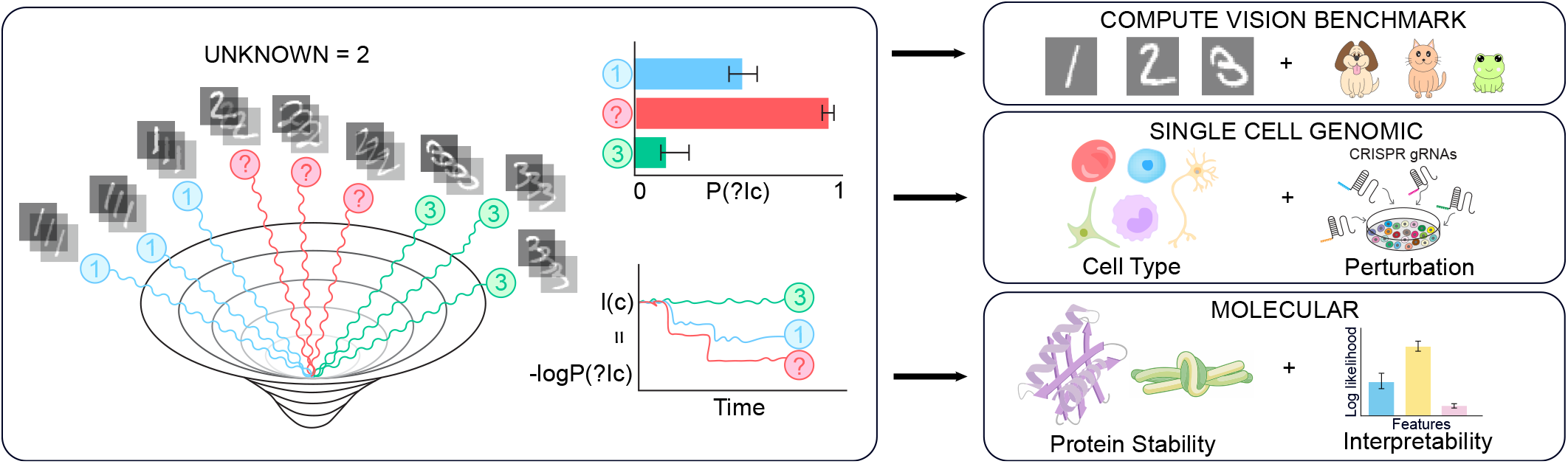
Generative classification from diffusion models with keeping score. Given a conditional diffusion model, the class log-likelihood of a new unseen sample (in this case ‘?=2’) can be inferred by noising the sample and calculating a time-dependent score-parameterized functional. We illustrate this new interpretable thermodynamic classification framework on computer vision tasks, problems from single cell genomics, and molecular/protein biophysics.

## RESULTS

In Box 1, we present an exact theory of thermodynamic classification with diffusion models, showing that for general diffusion processes, including Variance preserving (VP) and variance exploding (VE) diffusions, the likelihood *p*(*x*|*c*) can be expressed in terms of only a forward noising trajectory *x*_*t*_ with *x*_0_ = *x*, and the score function *s*_*θ*_(*x*_*t*_, *t, c*). With multiple noising trajectories, we can calculate error bars for the likelihood, and our results are interpretable, with the ability to prescribe a probability of successful classification for every input feature or degree of freedom in the problem. We demonstrate our approach on a range of problems from classical image datasets and single cell genomics, where we employ pretrained score functions, as well as an example from protein folding thermodynamics, training the score function to predict stability changes directly.

Keeping SCORE differs from discriminative classifiers because it interprets classification through the geometry of the underlying class-conditioned distributions rather than through learned decision boundaries. XGBoost and SHAP can identify which features push a prediction toward one class, but this explanation is post hoc and tied to the classifier output, not to an explicit probabilistic model of the data. In contrast, Keeping SCORE computes class likelihoods from a path functional of the conditional score field. The same likelihood used for prediction decomposes into coordinate-wise contributions, making attribution intrinsic to the inference procedure. Thus, Keeping SCORE provides generative, distribution-aware explanations, along with trajectory-based uncertainty, for why a sample is more compatible with one class manifold than another.

## Box 1.

**Score-Based Likelihood Theory and Classification**

### Score and forward diffusion

For class *c* and diffusion time *t*, the score is

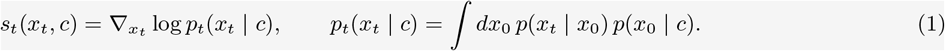

In continuous time, variance-preserving diffusion obeys

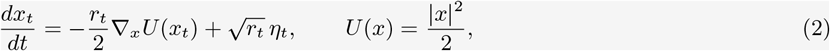

with ⟨*η*_*t*_⟩ = 0 and 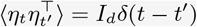. The associated Fokker–Planck equation is

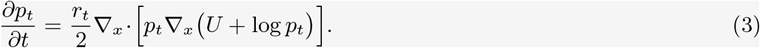

### Likelihood along a noising trajectory

For an individual stochastic path,

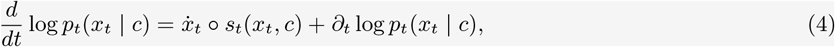

where ° denotes the Stratonovich product. Integrating and using Eq. (3) gives

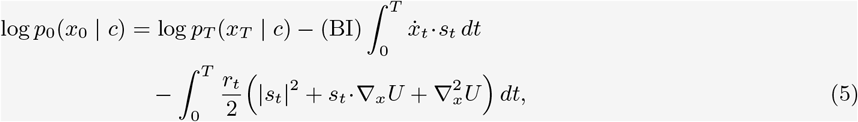

where *(BI)* denotes backward Itô evaluation, so score functions are evaluated at the end of the timestep for computational efficiency (SI Appendix 1, SI Appendix Fig. S1). For large *T, p*_*T*_ (*x*_*T*_ | *c*) = 𝒩 (*x*_*T*_ ; 0, *I*_*d*_), so this and other class-independent endpoint terms cancel in likelihood ratios. Thus

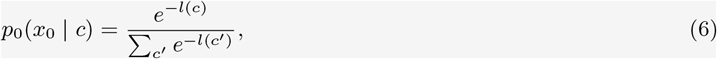

with

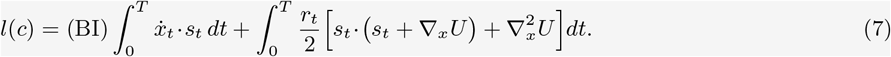

For VP diffusion, ∇_*x*_*U* = *x*, so 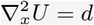 is constant in *c* and may be dropped for classification.

### Classification procedure

Given a new sample *x*_0_:

1. Generate a stochastic forward noising trajectory 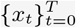 from Eq. (2).
2. For each candidate class *c*, evaluate the path functional *l*(*c*) from Eq. (7), then convert to class likelihoods *p*_0_(*x*_0_ | *c*) ∝ *e*^−*l*(*c*)^ and rank classes accordingly.
3. Repeat over independent noising trajectories initialized from the same *x*_0_ to estimate the mean and variance of *p*_0_(*x*_0_ | *c*), yielding uncertainty bars on class probabilities and ranks.

### Regression procedure

Given a new sample *x*_0_:

1. Generate a stochastic forward noising trajectory 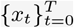 from Eq. (2).
2. Infer the prediction by optimizing *l*(*y*), a functional of the score neural network, with respect to *y*, i.e. *ŷ* = arg min_*y*_ *l*(*y*), or equivalently by gradient descent on *y*.
3. Repeat over independent noising trajectories initialized from the same *x*_0_ to estimate the mean and variance of *ŷ*, yielding uncertainty intervals on the prediction.

### Interpretability

Expanding in any orthogonal basis 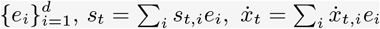 gives

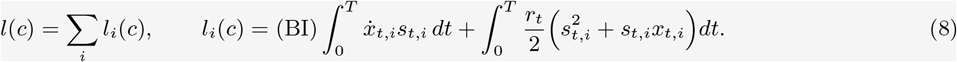

Each *l*_*i*_(*c*) measures the contribution of coordinate *i*—for example a gene, latent feature, or principal component—to the total class evidence.

## Classifying images with diffusion models

To benchmark our framework on a well-established vision task, we applied it to the MNIST dataset, which contains 70,000 28x28 pixel hand-written numbers, labeled 0-9 [23]. To allow for diffusion in a lower dimensional continuous space, we first trained a conditional convolutional variational autoencoder (VAE) with 10 latent dimensions. The images separate well by class in the latent space, as illustrated by UMAP (Fig. 2A). Next, we trained a denoising diffusion model on the latent space training points (6000 per class). Subsequently, we passed each of the test images (1000 for each of the 10 classes), through the VAE to get its position in latent space. From here, we noise the test points, integrating the log-likelihood for each of the 10 classes and calculating the posterior via soft-max. We show one such example, for a test point from the ‘5’ class (Fig. 2B, additional examples in SI Appendix, Fig. S2), where we averaged the posterior over 64 independent noising paths to perform uncertainty estimates. Overall, our diffusion based classification approach performs well on MNIST, with an AUPRC approaching 1 (Fig. 2C). To interpret which features are picked out as important for accurate digit classification, we calculate log-likelihoods for each of the ten latent space dimensions and linearize the decoder to connect each of these dimensions to the basic features of the problem, the pixels themselves. Such a pixel-wise attribution is depicted for a sample ‘5’ from the test set (Fig. 2D). Together, our framework illustrates the ability of generative models to quantify feature importance in image data, and could serve useful in the future to extract distinguishing features from medical imaging datasets, between healthy and diseased samples, for example.

**FIG. 2.**
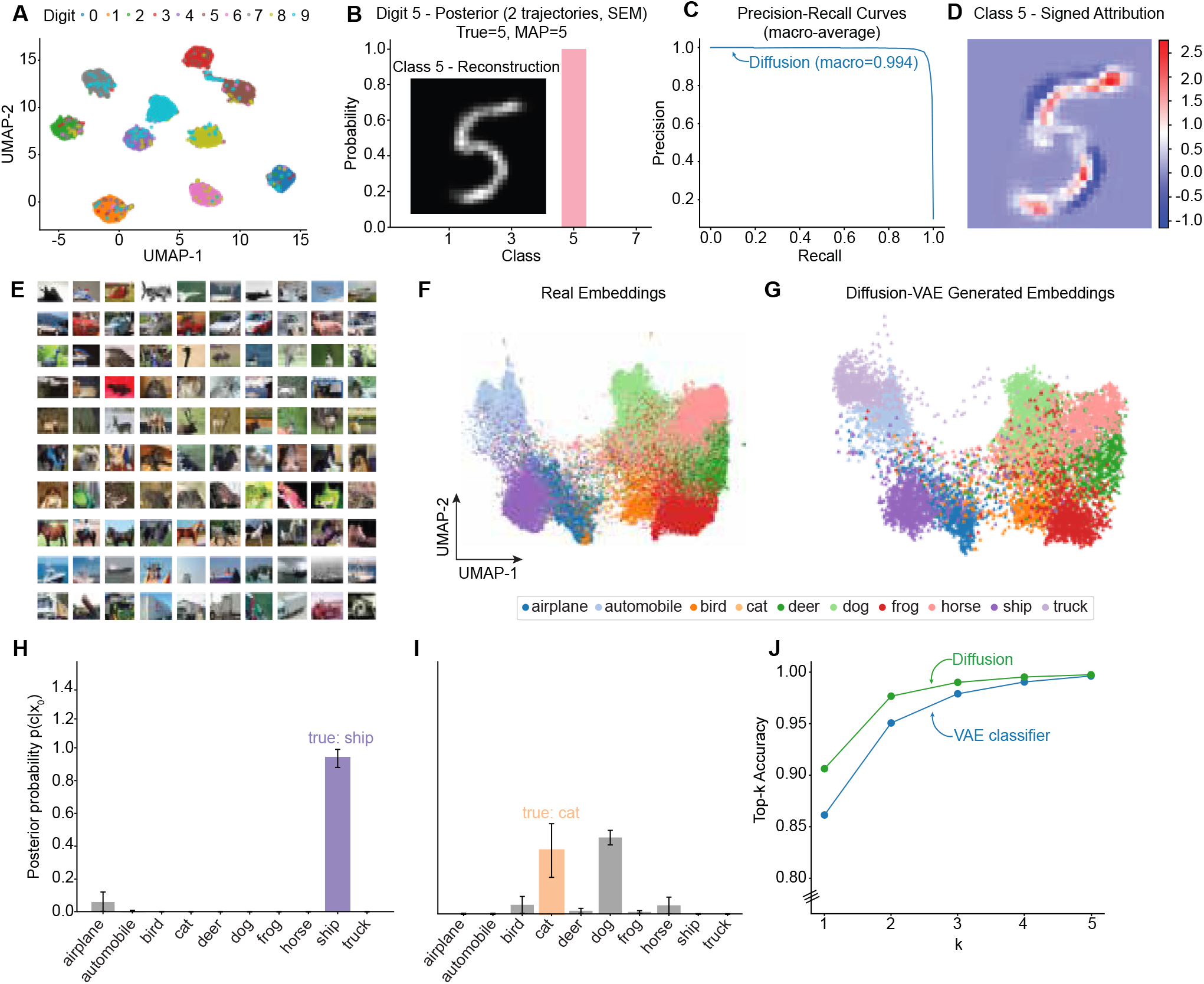
Diffusion classification for imaging data. MNIST: A) UMAP of test MNIST points in the latent space of a convolutional variational autoencoder. B) Posterior probability of a particular test point ‘5’, and reconstruction of the test point inset. C) Precision-recall curve for the test set. D) Pixel-wise pseudo-log-likelihoods for the reconstructed sample. CIFAR: E) Example images from the 10 classes in CIFAR10. F) UMAP of real reconstructed test embeddings, and G) UMAP of diffusion generated embeddings. H) Example posterior probability without a tie (overlapping error bars), and I) with a tie. J) Top-k accuracy reported for the diffusion classifier, the VAE classifier and tie aware diffusion classifier.

We next turned to the CIFAR10 computer vision benchmark, a dataset of 60,000 images spanning 10 object classes (airplanes, cats, dogs, ships, etc.) of size 32×32 pixels, each with 3 colors (Fig. 3E). CIFAR-10 provides a natural testbed for comparing generative and discriminative approaches because the classes have overlapping yet structured manifolds in pixel space. We first extracted embeddings from a pretrained ResNet-18 encoder to obtain compact representations of each image. These embeddings were then used to train a variational autoencoder (VAE), providing a smooth lower-dimensional latent manifold on which we trained the diffusion model. We illustrate concordance between the original data and that generated by the diffusion process via UMAP of the original 512 dimensional test embeddings, alongside those from the diffusion model (Fig. 2F, G). By evaluating class likelihoods of the 10,000 previously unseen test images along diffusion trajectories in the VAE latent space, we recovered the true class with an accuracy comparable to the built-in Multi-layer Perceptron (MLP) classifier with which the VAE latent space was shaped. Crucially, we were able to account for uncertainty in the class probabilities by averaging them over just two independent noising trajectories (Fig. 2H, I and SI Appendix, Fig. S3). Keeping SCORE achieved higher scores than the VAE neural network classifier used to shape the latent space (Fig. 2J).

**FIG. 3.**
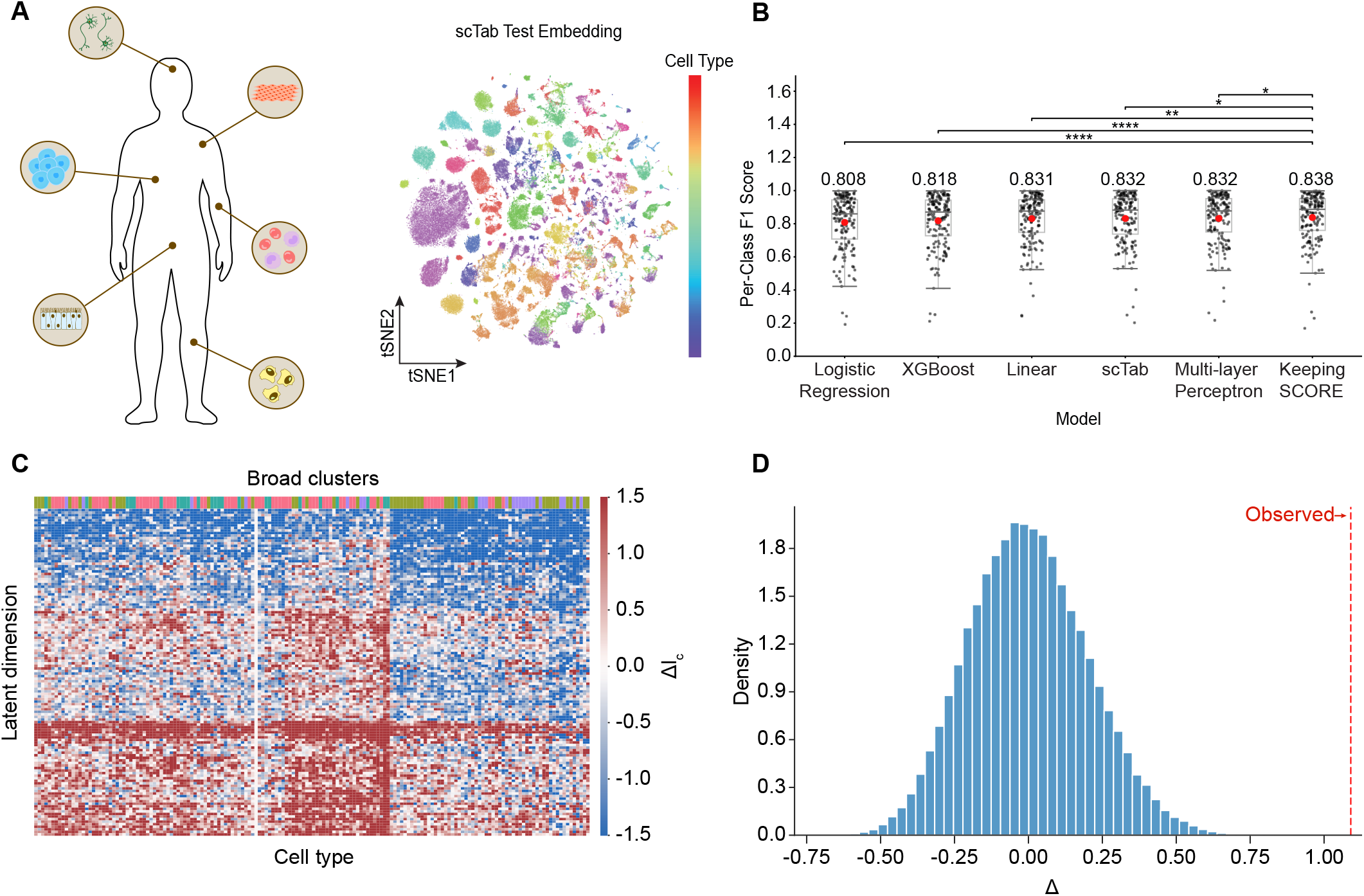
Diffusion classification for single-cell genomics. A) Cell types from primary tissue in the body (left) and t-distributed stochastic neighbor embedding (tSNE) plot of scTab cell type atlas test embedding [25] (right). B) The cell type classification results from various machine-learning algorithms (from left to right: logistic regression, XGBoost, linear model, scTab, MLP, and Keeping SCORE). Each small black particle represents the per-class F1 score across 164 cell types. Box plot embodies median (middle line), interquartile range (hinge), and 1.5 times of interquartile range (whisker). The number above the boxplot represents the macro-average F1-score (also marked as a large red dot on each box plot). The Keeping SCORE macro-average F1-score was adjusted based on the tie analysis (see Materials and Methods). The original macro-average F1-score for Keeping SCORE without tie awareness is 0.828. Statistical significance is calculated based on Wilcoxon signed-rank test p-value adjusted with false discovery rate (FDR). ns: *p* ≥ 0.05, * : *p <* 0.05, ** : *p <* 0.01, * * * : *p <* 0.001, * * ** : *p <* 0.0001. C) Latent space log-likelihood attribution Δ*l*_*c*_ = *l*_*c*_ − *l*_*ref*_ heat map as a function of cell type, clustered over latent dimensions and cell type and labeled by broad clusters (defined by KNN clustering of latent space cell type class centroids, K=5). Attributions are measured with respect to a fixed reference cell type (alveolar type 1 fibroblast cell). Exact p-value from the permutation analysis with 100,000 permutation, *p <* 10^−5^ (two-sided test). D) Average pairwise distance between two latent space attribution-vectors: one representing the pairwise distance between two cell types across broad (KNN) clusters, and one from within broad clusters, Δ = ⟨*δr*⟩ _*between*_ − ⟨*δr*⟩ _*within*_. The observed value of Δ is shown in red (observed value = 1.138), and the distribution is over random null samples where the broad clusters have been randomly distributed across cell type.

## Classifying cell type from gene expression

Multicellular tissues comprise diverse cell types, each defined by distinct transcriptional programs that give rise to specialized functions [26, 27] (Fig. 4A). In single-cell transcriptomics, cell types are typically annotated using combinations of known marker genes. This approach is interpretable but limited by prior knowledge and by the nonlinear overlap between transcriptional states. To test whether our approach could learn these distinctions directly from data geometry, we used neural network embeddings of 164 unique cell types derived from the scTab [25] harmonized single-cell atlas (see Materials and Methods). We trained a diffusion model on a scTab embedding [25] and evaluated performance on the subsampled held-out test classes (49,046 datapoints (cells)) (see Fig. 3A and SI Appendix, Fig. S4). By comparing diffusion-based likelihoods, we are able to classify unseen expression profiles into their most probable cell type. When overlapping error bars were taken into account (see Materials and Methods), our model shows modest improvement over both simple (logistic regression, linear perceptron) and state-of-the-art (XGBoost, scTab, multi-layer perceptron) machine-learning models [25] (Fig. 3B).

**FIG. 4.**
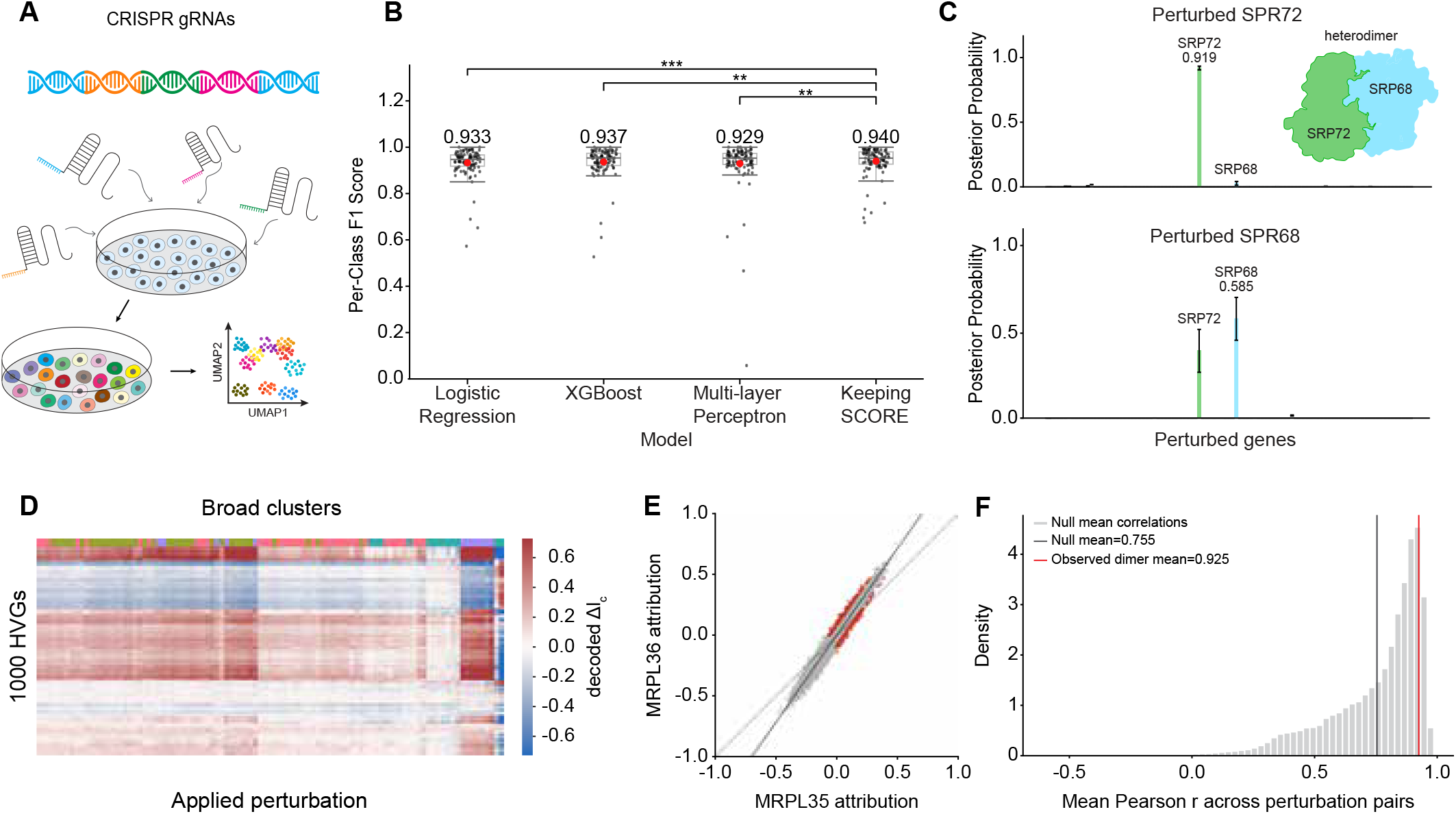
Diffusion classification for single-cell genomics. A) CRISPR gene perturbations induce different phenotypes in cellular populations. B) The genetic perturbation classification results from various machine-learning methods (from the left-side: logistic regression, XGBoost, MLP, and Keeping SCORE). Each small black particle represents the per-class F1 score across 100 genetic perturbation types. Box plot embodies median (middle line), interquartile range (hinge), and 1.5 times of interquartile range (whisker). The number above the boxplot represents the macro-average F1-score (also marked as a large red dot on each box plot). The Keeping SCORE macro-average F1-score was adjusted based on the tie analysis (see Materials and Methods). The original macro-average F1-score for Keeping SCORE without tie awareness is 0.932. Statistical significance is calculated based on Wilcoxon signed-rank test p-value adjusted with false discovery rate (FDR). ns: *p* ≥ 0.05, * : *p <* 0.05, ** : *p <* 0.01, * * * : *p <* 0.001, * * ** : *p <* 0.0001. C) The plots show the posterior probability of two different genetic perturbations: SRP72 (upper) and SRP68 (lower). In the upper figure, SRP72 is mostly correctly mapped to SRP72 (0.919 ± 0.015) where the second prediction is SRP92 (0.027 ± 0.015). However, in the lower figure, about half of SRP68 (0.585 ± 0.126) is mapped to SRP72 (0.396 ± 0.126). D) Gene space (original data dimensionality) log-likelihood attribution Δ*l*_*c*_ = *l*_*c*_ − *l*_*ref*_ heatmap as a function of perturbed gene type, clustered over the gene space dimensions and perturbation type and labeled by broad clusters (defined by KNN clustering of gene space perturbation type class centroids, K=6). Attributions are measured with respect to a fixed control cell centroid (the mean expression level of the decoded control cells). E) MRPL35 attribution vs MRPL36 attribution in the decoded gene space. The attributions are normalized by division of the highest attribution value. Grey dots present the attribution of the target perturbation to that gene. Red dots indicate genes with positive attributions for either MRPL35 or MRPL36 with magnitude greater than two standard deviations from the mean (black line). F) Distribution of Pearson correlations between pairs of randomly chosen perturbed genes. The mean of the distribution is shown as a grey dashed line. The black dashed vertical line indicates mean of Pearson correlation between the three pairs of naturally occurring as a component of complexes in the perturbation data: MRPL35-MRPL36, SRP68-SRP72, and UPF1-UPF2.

To determine whether biologically meaningful structure organizes the diffusion likelihood rather than purely discriminative decision boundaries, we examined how individual latent dimensions contribute to the class log-likelihood. For each cell-type centroid, we computed a coordinate-wise attribution vector Δ*l*_*c*_ = *l*_*c*_ − *l*_ref_, measuring the contribution of each latent dimension to the likelihood of that cell type relative to a fixed reference cell type. We then clustered cell types using a KNN graph constructed from their latent-space representations, producing broad cell-type groups that summarize coarse transcriptional organization (Materials and Methods, SI Appendix Fig. S5). When we clustered the resulting attribution matrix across both cell types and latent dimensions, cell types belonging to the same broad group formed largely contiguous blocks with shared attribution structure (Fig. 3C). This pattern suggests that the latent degrees of freedom that Keeping SCORE learns are not arbitrary classifier features, but encode recurrent transcriptional axes that distinguish biologically related families of cell types. To quantify this effect, we compared distances between attribution vectors for cell types in different broad clusters to those within the same broad cluster, defining Δ = ⟨*δr*⟩ _between_ − ⟨*δr*⟩ _within_. The observed value of Δ was substantially larger than expected under random reassignment of broad-cluster labels, indicating that cell types within the same coarse biological group share significantly more similar likelihood-attribution profiles than unrelated cell types (Fig. 3D, *p* − *value <* 10^−5^). When testing on the mean gene expression of each cell type (164 datapoints), we observed that evaluating with significantly fewer steps did not dramatically affect classification performance (Fig. S6, Materials and Methods). This idea may be expanded to knowledge distillation [28] to reduce the computational cost. Together, these results suggest that thermodynamic class structure can recapitulate biologically meaningful cell-type hierarchies. In the future, this framework could both identify interpretable marker gene sets and annotate previously unseen cells.

## Classifying genetic perturbation

Understanding the intricate relationship between genetic features and observable phenotypes is central to genomics research. Modern functional perturbation screens, such as those using CRISPR interference or activation (CRISPRi/a) to suppress or activate the gene(s) of interest, systematically alter gene expression to reveal how individual gene perturbations and their combinations influence cellular states and macroscopic phenotypes. Recent methods, such as Perturb-seq [29] or CROP-seq [30], enable massively parallel genotype-to-phenotype screening by introducing thousands of CRISPR-based perturbations in a single experiment, while simultaneously capturing molecular readouts at single-cell resolution (Fig. 4A). While these datasets provide a powerful window into measurable causal gene–function relationships, a central question remains unresolved: which perturbations drive which phenotypic transitions, and through what combinations of gene-level effects [9]? Resolving this mapping is critical to understanding and ultimately mitigating complex disease phenotypes driven by dysregulated gene networks. To test our framework in this context, we used VAE to create embeddings of single-cell transcriptomic profiles from 100 unique gene perturbations (Fig. 4B inset, Materials and Methods). We trained a diffusion model on a subset (67,808 datapoints) of these perturbations and reserved the remainder for testing (8,477 datapoints) (SI Appendix, Fig. S7). Next, we computed the mean embedding of each test perturbation and classified it based on its diffusion likelihood (Fig. 4E, Materials and Methods). Again, we found our approach to be favorable in classification score to other baselines: logistic regression, XGBoost, and a multi-layer perceptron). Direct likelihood evaluation allowed us to gauge how well a generative model could capture the global organization of perturbation-induced expression changes.

Beyond aggregate classification performance, the likelihood-based formulation provided biologically meaningful uncertainty estimates and gene-level interpretability. For each held-out perturbation, Keeping SCORE returned a full posterior distribution over candidate perturbed genes rather than a single class label. Notably, some ambiguities in this posterior reflected real biology. For example, SRP72 was assigned confidently to the correct perturbation (Fig. 4C, upper plot), whereas SRP68 showed substantial posterior probability split between SRP68 and SRP72 (Fig. 4C, lower plot). We were able to biologically rationalize this ambiguity: SRP68 and SRP72 form a heterodimeric subcomplex of the signal recognition particle S domain [31], so perturbing one member can disrupt the same molecular module and may lead to shared pheotypic changes. [32, 33]. More critically, the posterior distributions were asymmetric: SRP68 confusion bled into SRP72 but not vice versa. We sought to resolve this directly. SRP68 directly mediates binding of the SRP68/72 heterodimer to 7S RNA at the three-way junction region [34–36], whereas SRP72 stabilizes the interaction more indirectly through electrostatic contacts [35]. Perturbation of SRP68 may therefore produce transcriptional changes in SRP72, without the reciprocal effect. We provide additional symmetric and asymmetric cases in SI Appendix, Fig. S8. This uncertainty is particularly informative in perturbation biology: it points to shared complexes, compensatory structure, or functionally coupled genes, and it arises naturally from Keeping SCORE’s posterior estimation over perturbations. Standard discriminative or flow-based classifiers would require additional calibration or ensembling to produce comparable estimates.

We next asked whether the likelihoods were organized by biologically meaningful transcriptional structure (Material and Methods; latent embedding visualized in SI Appendix, Fig. S9). To do so, we decomposed the class log-likelihood relative to a control reference into decoded gene-space contributions and clustered the resulting attribution matrix across both perturbations and genes. The resulting heatmap showed block structure aligned with broad perturbation clusters defined independently from perturbation centroids, indicating that the likelihood contributions reflected shared transcriptional programs among related perturbations rather than unstructured classifier weights (Fig. 4D). Note that both Keeping SCORE and tree-based classifiers like XGBoost can yield gene-level attribution maps, but they differ in what they explain: Keeping SCORE attributes genes to the perturbation-conditioned likelihood before classification, whereas tree-based classifiers explain a learned decision boundary. It should also be noted that both methods (Keeping SCORE and XGBoost) extrapolate from latent to gene-level features with the same linear decoder. Together, our analysis suggests that the likelihood attributions could be capturing coordinated biological structure in the perturbation response. For example, the mitochondrial ribosomal proteins MRPL35 and MRPL36 show highly similar gene-level attribution profiles, with the strongest positively attributed genes lying close to the diagonal. This indicates that such perturbations are based on overlapping transcriptional evidence (Fig. 4E). More generally, naturally occurring protein complexes, including MRPL35–MRPL36, SRP68–SRP72, and UPF1–UPF2, exhibited higher mean attribution-profile correlations than randomly chosen perturbation pairs (Fig. 4F). Although this enrichment was not statistically significant given the small number of annotated pairs, it supports the interpretation that Keeping SCORE assigns similar likelihood structure to perturbations whose gene products participate in shared physical or functional complexes. In the future, gene-space likelihood attributions may provide a useful route toward perturbation prediction beyond classification accuracy alone, by identifying when distinct perturbations act through shared transcriptional programs and by prioritizing candidate functional relationships, complexes, or compensatory perturbations for experimental follow-up.

## Predicting how mutations change protein stability

Protein stability reflects the balance between folded and unfolded conformations, and single amino acid mutations can shift this equilibrium by an amount quantified by ΔΔ*G*—the change in folding free energy relative to the wild type, where positive ΔΔ*G* indicates destabilization, corresponding to a larger fraction of unfolded proteins in solution (Fig. 5A)[37–39]. Accurately predicting ΔΔ*G* from sequence is vital for understanding misfolding diseases, engineering more stable proteins, and identifying pathogenic susceptibilities in known structures. For this task, we rely solely on the local linear amino acid sequence surrounding each mutation, with no explicit structural or chemical information. In this setting, our approach performs regression rather than classification, treating ΔΔ*G* as a continuous conditioning variable in the diffusion model (Fig. 5B). We infer stability changes by performing direct gradient descent on the diffusion likelihood with respect to ΔΔ*G*, finding the value that maximizes consistency with the observed experimentally measured value. This optimization can be applied to pretrained diffusion models or incorporated directly into training by differentiating through the gradient descent step itself, minimizing the deviation between predicted and true ΔΔ*G* values. We use a combined loss function

**FIG. 5.**
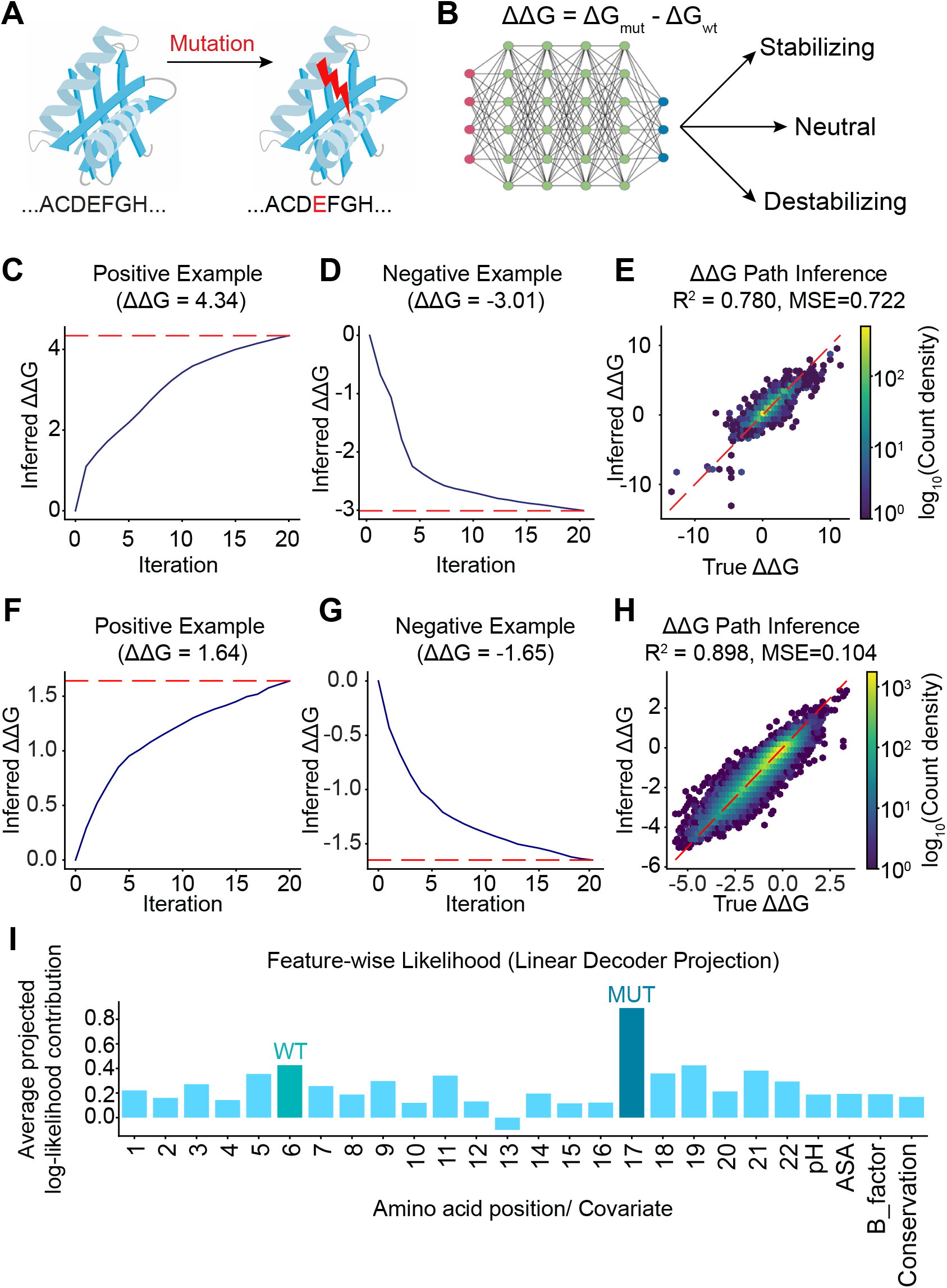
A) Protein with a single amino acid mutation. B) Regressing protein stability change ΔΔ*G* upon mutation with score based diffusion models. C), D) Gradient descent for inferred stability change over iterations; true stability change shown as red dashed line. E) Inferred and true ΔΔ*G* counts for all points in the test set. F)-H) The same analysis as C)-E) but with the mega-scale protein stability dataset, split by mutation. I) Average (over two independent proteins and mutations) log-likelihood for input model features: amino acid number relative to mutational origin, and experimental covariates. Location of WT and mutated amino acids shown in dark blue.

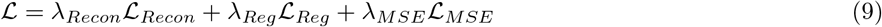

where ℒ_*Recon*_ is the reconstruction loss, ℒ_*Reg*_ the classical regression head, and ℒ_*MSE*_ is the mean-squared-error between the real stability change, and that predicted, *ŷ*_*i*_ = ΔΔ*G* = argmax_*y*_ *l*(*s*(*y*)), by gradient descent:

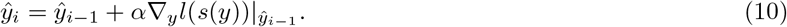

Here the *λ* parameters are weighting constants (Materials and Methods).

We consider a neighborhood of ten amino acids (five on each side) around the mutated cite. As input features, we concatenate the pre-mutation amino acid sequence with the mutated sequence as well as four additional covariates: indicating experimental pH, evolutionary conservation of the amino acid, solvent accessible surface area (ASA), and *B*_*factor*_ scores, the last of which quantifies how much an atom’s electron density is “smeared out” due to thermal motion, static disorder, or uncertainty in atom position during structure refinement. We use the FireProtDB [40] database with a total of 31, 223 measured examples, which we split 70/15/15 into training, validation, and test sets.

We train the full joint loss for 2500 epochs with a small number of diffusion steps *T* = 2. Once trained, this small diffusion time allows for rapid inference. We show two examples of the gradient descent process on the log-likelihood for both highly stabilizing and destabilizing test set mutations, illustrating convergence to the correct answer in the number of gradient descent steps the model was trained with (Fig. 5C,D). A histogram of the true and inferred ΔΔ*G* values shows a high *R*^2^ ≈ 0.78 with a MSE of about 0.72 [kcal*/*mol] (Fig. 5E). We observe similarly high performance on an independent mega-scale protein stability dataset (Fig. 5F-H, SI Appendix Fig. S10) [41], trained with identical architecture but distinct proteins, mutations, and covariates—indicating robustness to experimental specifics. Our performance, as measured by Root Mean Squared Error (RMSE), on two kinds of data splitting for training/testing (partition by mutation, *RMSE* = 0.322 [kcal*/*mol], or partition by protein, *RMSE* = 0.713 [kcal*/*mol], *Pearson* − *r* = 0.761) matches another recent structure-informed transfer learning study which split the data by protein and achieved *RMSE* = 0.708 [kcal*/*mol], *r* = 0.754 [42].

We train the VAE with a linear decoder, so that we can directly attribute likelihood values from the non-linear diffusion classification in the three dimensional latent space to each amino acid and covariate, both on average and on the level of individual proteins/mutations. We find that for a particular pair of proteins, the pre- and post-mutated amino acid are the features that most effect classification at the given inferred stability change (See Fig. 5I and SI Appendix, Fig. S11). These results demonstrate that even a shallow diffusion process, coupled with an interpretable latent representation, can extract the local biochemical determinants of protein stability and provide residue-level mechanistic insight as to how important the local neighborhood around a mutation, and the mutation itself, is to driving measured stability change.

## DISCUSSION

Here we developed a novel framework called Keeping SCORE that repurposes diffusion models as inference engines for interpretable, uncertainty-aware classification and regression. By computing exact class likelihoods from a path observable along forward noising trajectories, our approach requires no approximations, architectural modifications, or retraining of existing diffusion models. We demonstrated that the conditional score function naturally encodes distributional structure, enabling both accurate prediction and decomposition of classification decisions into feature-specific contributions. Applied across computer vision benchmarks, single-cell genomics, and protein stability prediction, Keeping SCORE reliably quantified posterior probabilities, attributed predictions to biologically or physically meaningful degrees of freedom, and provided error bars through trajectory averaging. Our work establishes diffusion models as general-purpose Bayesian inference machines that unify generation with interpretable discrimination for genomics.

Recent efforts to extract classification signals from diffusion models have relied on approximating likelihoods through the evidence lower bound (ELBO) [16, 17], which provides a tractable but inexact estimate of the log-probability. While these approaches demonstrate the promise of diffusion-based classification, they sacrifice exactness and lack mechanisms for attributing predictions to specific input features. More broadly, several studies have connected path-integral formulations to diffusion model likelihoods—through perturbative expansions that noted exact computation along stochastic paths remained open [18], stochastic interpolant estimators for generative design rather than classification [19, 20], and fluctuation theorems from statistical physics applied to sampling and free energy estimation [21, 22]. In contrast, our approach computes likelihoods exactly via path integrals, yielding predictions that decompose naturally into interpretable coordinates. These aspects position our framework as particularly valuable for medical genomics domains like patient stratification and disease classification, where both uncertainty quantification and mechanistic insight are essential. Deeper connections between statistical physics and machine learning may open new avenues for principled, physics-grounded model design in genomics. A particularly relevant study recently showed that free energy differences can be computed efficiently with diffusion models [43], which differs from the present work in that we consider classification and regression tasks, and work with the widely used VP diffusion instead of alchemical potentials.

A complementary perspective comes from recent work connecting diffusion models to hierarchical decision trees. Tree-based classifiers such as XGBoost achieve strong performance by recursively partitioning feature space into increasingly refined regions, producing an explicit discrete hierarchy of decisions. Diffusion models appear to learn an analogous hierarchy in continuous time: as noise is varied, high-level semantic variables such as class can undergo sharp transitions, while lower-level features evolve more gradually and may persist across class changes [44]. This view is supported by recent analyses showing that diffusion dynamics probe hierarchical and compositional structure in data [45, 46], and by theoretical work relating decision-tree coarse-graining to probability-flow dynamics [47]. In this sense, Keeping SCORE can be viewed as a generative, continuous-time analogue of tree-based classification: rather than assigning a cell or perturbation profile to leaves of a learned decision partition, it evaluates how likely it is under each class-conditioned generative trajectory. This distinction matters for interpretability and uncertainty estimation. Where XGBoost explains predictions through splits or feature gains attributed to a discriminative boundary, Keeping SCORE decomposes an explicit likelihood ratio into coordinate-wise contributions along the diffusion path, exposing hierarchical structure as Bayesian evidence, posterior uncertainty, and path-wise (e.g., gene-level) feature attribution. Our approach has certain limitations as well: primarily, the computational cost of evaluating diffusion models, which often requires dimensionality reduction through autoencoders or pretrained embeddings for tractable inference. Such preprocessing steps may discard information or introduce artifacts that affect predictions. Future work should explore efficient score architectures as well as adaptive sampling strategies (analogous to tau-leaping in stochastic simulations [48]) and model distillation that concentrate computation on informative diffusion timesteps. We also expect future adaptations of single-step Flow Maps [49] to perform interpretable, uncertainty aware classification without requiring dimensionality reduction to a low rank latent space. Additionally, our current implementation assumes the score function is well-trained and accurately captures the true distribution; poorly trained diffusion models will propagate errors into classification decisions. We note, however, that our approach is naturally amenable to Monte Carlo sampling the posterior distribution. This means that Keeping SCORE can also apply well to combinatorial problems, like multigene perturbation prediction, where the number of classes grows exponentially.

Looking ahead, Keeping SCORE opens promising avenues for extension. Our framework extends to genomics and protein design applications, from identifying interpretable marker gene sets for novel cell types and mutational landscape for protein-ligand binding, to predicting responses to combinatorial perturbations, and to performing Bayesian model selection among competing generative hypotheses of cellular state. Extending to temporal data, such as classifying developmental trajectories or predicting cellular state transitions from time-series expression, could capture tipping points in dynamics. Scaling to multigene perturbations remains crucial for translating our approach to complex disease modeling. Overall, our interpretable generative classification framework is poised for mechanism discovery and control across genomics and molecular biology.

## MATERIALS AND METHODS

### The path likelihood *l*(*c*) as a ratio of forward and effective backward path ensembles

Here we demonstrate the relationship

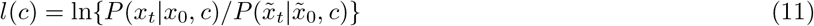

where 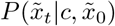 is the 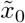-conditioned path probability of the time-reversed trajectory 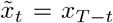 under the time-reversed Langevin equation

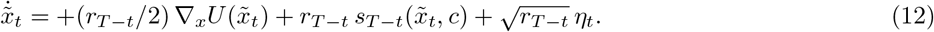

Eq. (11) is similar to that of *heat dissipation* in stochastic thermodynamics, though strictly speaking, dissipation is quantified by path ratios under the same ‘protocol’, without the additional score function [50]. Here, however, the reverse process (12) is different from the forward process (Box 1), as an additional force (the score function) appears. In this way, *l*(*c*) is the heat dissipation corresponding to the asymmetric-protocol work 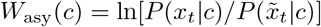 studied in [22].

The log path probability of the forward process is given by ln *P* (*x*_*t*_|*x*_0_, *c*) = −*S*[*x*_*t*_|*c*] + const, where

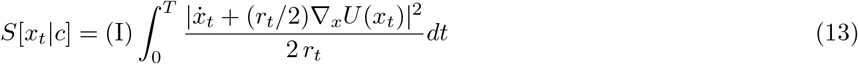

is the appropriate Onsager-Machlup path action, and (I) denotes a forward Itô integral [51]. For the reverse process, ln 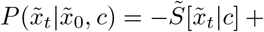 const, where

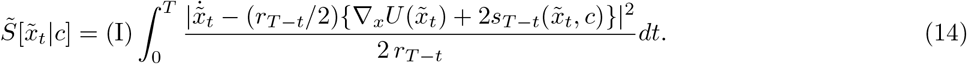

In order to relate 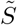 to *S*, one must perform the transformation *t* → *T* − *t*, which yields

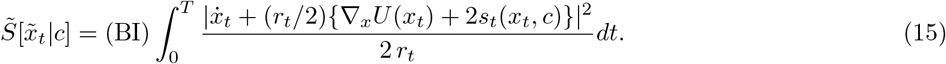

Finally, we can subtract the two path actions 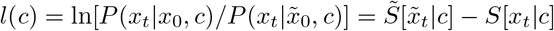 to get

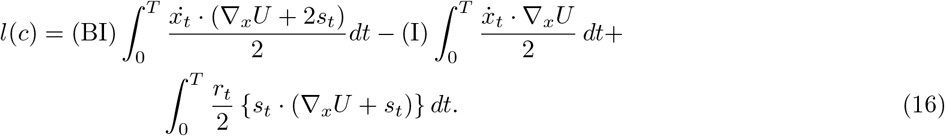

Now, there is a choice to be made as to whether to convert the three integrals into a single Itô, Stratonovich, or backwards Itô integral (see Appendix A.2 of [22]), yielding the three separate but equivalent expressions:

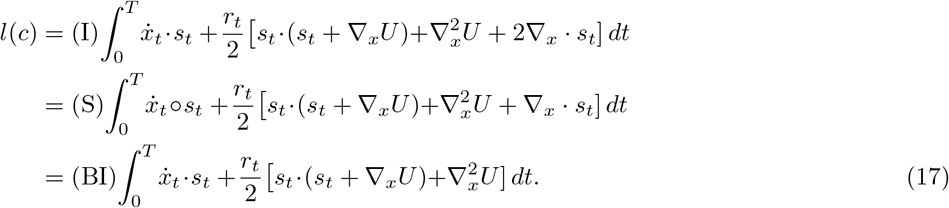

It is computationally expensive to calculate ∇_*x*_ · *s*_*t*_, but conveniently it does not have to be calculated in the backwards Itô integral (see SI Appendix 1, Fig. S1) .

### Classifying natural images with diffusion models

We trained a convolutional variational autoencoder (VAE) on MNIST digits using extensive online augmentation (random rotations, affine transformations, translations, perspective distortions, and scaling), with a separate validation set using only tensor conversion. The encoder consists of two stride-2 convolutional layers (1 → 32 → 64 channels with 4 × 4 kernels, BatchNorm, ReLU), flattened and mapped to a 2D latent Gaussian via fully connected layers predicting the mean and log-variance. Latent samples are drawn with the reparameterization trick. The decoder inverts this architecture, expanding the latent vector back to a 64 × 7 × 7 feature map followed by two transposed-convolutional blocks (64 → 32 → 1) ending in a Sigmoid pixel likelihood. A linear classifier operating directly on the latent vector predicts the digit label. The loss function combines a weighted binary cross-entropy reconstruction term (scaled by 1000), KL divergence between the approximate posterior and 𝒩 (0, *I*), and a latent classification cross-entropy term (scaled by 200). The model was trained for 600 epochs with large batches (3000 samples) using the Adam optimizer (learning rate 10^−3^), and after each epoch all validation images were encoded to collect latent embeddings for downstream analysis.

To model a class-conditional structure in the learned 2D latent space, we trained a conditional denoising diffusion model directly on the VAE latent codes. For each latent–label pair (*z*_0_, *y*), a random timestep *t* ∈ {0, …, *T* − 1} with *T* = 1000 was sampled, and a corrupted latent *z*_*t*_ was generated. A class-conditional denoiser predicted *ϵ* from (*z*_*t*_, *t, y*), using separate embeddings for time and class concatenated with *z*_*t*_ and passed through a three-layer fully connected network (512 units, GELU). The training objective was mean-squared error between predicted and true noise, optimized with Adam (learning rate 10^−3^) for 1000 epochs using large batches (5000). The best-performing denoiser was selected by lowest validation loss. Together, the latent VAE and the conditional diffusion model enable class-guided generation of novel digit images by sampling *z* via diffusion dynamics and decoding through the VAE decoder.

We trained a joint variational autoencoder and diffusion-based model on fixed ResNet-18 embeddings of the CIFAR-10 dataset. Precomputed embeddings containing 512-dimensional feature vectors for each training image and corresponding class labels were loaded. Prior to model training, all embeddings were standardized using a global mean–variance transform, and the resulting scaling statistics were saved for later use in downstream inference. We constructed a Progressive VAE with a 64-dimensional latent representation, consisting of a two-layer MLP encoder that outputs the mean and log-variance of the approximate posterior, and a symmetric MLP decoder that reconstructs the original embedding. A lightweight classifier was attached to the latent space to encourage class-separable representations. The VAE, classifier, and conditional diffusion model were trained jointly. For the diffusion component, we used a cosine *β*-schedule with *T* = 2500 forward noising steps, and trained a conditional denoiser on of the latent space. The denoiser receives the noised latent variable *z*_*t*_, the diffusion timestep *t*, and a learned class embedding, and predicts the forward noise using a residual MLP with sinusoidal Fourier time features.

Training was performed end-to-end with an AdamW optimizer over 250 epochs, using a batch size of 256 and weight decay of 10^−8^. The total loss consists of three terms: the VAE reconstruction plus KL regularization term, the cross-entropy classification loss, and the diffusion noise-prediction loss. The KL contribution was down-weighted (KL weight = 10^−6^) to stabilize optimization in the high-dimensional embedding space, while the classifier and diffusion losses were weighted equally (*λ*_cls_ = *λ*_diff_ = 1). The best-performing model checkpoint (lowest total loss) was saved and used for evaluation. This joint architecture enables simultaneous learning of a structured latent space, class-conditional structure, and diffusion dynamics directly over pretrained image embeddings.

### Classifying cell type from gene expression

The performance of each model was evaluated on the ability to label the cell type correctly on the single-cell transcriptional profile embeddings. These embeddings were obtained from scTab [25], a TabNet-based [52] state-of-the-art cell type classifier. According to the literature, the original ATLAS dataset contains approximately 22 million cells which were sampled from 56 different tissues from 5,052 donors, resulting 164 unique cell types. This ATLAS dataset was divided into a training set with 15,240,192 cells, a validation set with 3,500,032 cells, and a test set with 3,448,832 cells. These datasets were passed to the scTab feature transformer to obtain each 128-dimensional embedding, consistent with the experimental design of scTab (Table I). The embeddings were loaded following the original protocol: https://github.com/theislab/scTab. For benchmarking, the test embedding was downsampled to a maximum of 300 samples for each cell type class with a fixed seed of 42, resulting 49,046 datapoints. All of the listed models were trained, validated, tested, and benchmarked with the obtained embeddings. The Python environment setting followed the scTab guidelines with Python 3.8.

**TABLE I.** Cell type embedding splits and dimensions.

| Train set | Validation set | Test set |
| --- | --- | --- |
| 15,240,192 × 128 | 3,500,032 × 128 | 3,448,832 × 128 |

### Cell type classification benchmark

#### 1. Keeping SCORE

Keeping SCORE for the cell type classification task was built with a simple conditional diffusion model. The model hyperparameters are listed in Table S1. The model utilized the sigmoid beta schedule as it was effective for denoising the scRNA-seq dataset [53, 54]. The conditional denoiser was built with two hidden layers, each with 512 dimensions. Class labels were encoded through a learned 64-dimensional encoding vector. Timestep of 1000 was encoded via a two-layer MLP based on the GELU activation function. The diffusion model was trained based on the PyTorch Lightning module with the suggested learning rate ≈ 0.0021. During the training process, the diffusion model was validated by the validation embedding. The training was early-stopped after 313 epochs. The fully trained model was passed to Keeping SCORE with 4 noising paths. Keeping SCORE was run on GPU with vectorized and reshaped loops for the computational efficiency. The output from the diffusion model network was directly used as the score function value. Keeping SCORE’s structure utilized anthitetic pairing, meaning that for every noise *ϵ*, the paired − *ϵ* was included to improve Monte Carlo estimation stability. The uncertainty of noise path was reflected in the analysis as error bars. For the uncertainty aware analysis, we considered model’s prediction was correct when the highest classified class error bar (the standard error of each noise path) contains the true class mean.

For the timestep analysis, the same diffusion model was utilized with downsampling the timestep to 50, 100, 250, 500, 600, 700, 800, and 1000 in the Keeping SCORE analysis. Unlike a linear beta schedule, a sigmoid beta schedule timestep cannot be correctly downsampled with a linear sampling. Hence, we downsampled diffusion model sigmoid beta schedule with the variance-based subsampling instead of directly subsampling from the original timestep domain (also explained in SI Appendix 2).

#### 2. scTab

scTab model dataloader was customized to train on embeddings directly and bypass all the preprocessing steps. All the other training hyperparameters and model structure were kept the same as described in the original scTab model. The model was not fine-tuned and trained with the train embedding with the same PyTorch Lightning parameters as described in https://github.com/theislab/scTab/blob/devel/notebooks/training/train_tabnet.ipynb. The learning rate was set to ≈ 0.01995 by the Lightning. The model was evaluated on the subsampled test embedding.

#### 3. Multi-layer Perceptron

The MLP model from scTab was directly used for the cell type classification task https://github.com/theislab/scTab/blob/devel/notebooks/training/train_mlp.ipynb. The model was re-trained on the full train embedding loaded by the PyTorch Lightning module with the most optimum learning rate found by the learning rate finder (learning rate ≈ 0.0302). The validation loss was calculated based on the validation embedding. The model was evaluated on the subsampled test embedding.

#### 4. XGBoost

GPU-based XGBoost (version 1.6.2) was used for the cell type classification task. The XGBoost model was trained on the full train embedding loaded without any additional subsampling or modification. The validation loss was calculated based on the validation embedding. Each cell type class was weighted according to the class weights described in scTab. The model structure and training details can be found in Table S2. The model was evaluated on the subsampled test embedding.

#### 5. Logistic Regression

SGDClassifier from Scikit-learn (version 0.24.2) was used for the cell type classification task in lieu of CellTypist [55]. The model was trained on CPU with the full train embedding loaded without any additional subsampling or modification. The validation loss was calculated based on the validation embedding. The model structure and training details can be found in Table S3. The model was evaluated on the subsampled test embedding.

#### 6. Linear Model

The linear model from scTab was directly used for the cell type classification task https://github.com/theislab/scTab/blob/devel/cellnet/models.py. The model was re-trained on the full train embedding loaded by the PyTorch Lightning module with the most optimum learning rate found by the learning rate finder (optimum learning rate ≈ 0.01995). The data loading process was modified to load the embedding. During the training, validation loss was calculated based on the the validation embedding. The model was evaluated on the subsampled test embedding.

### Computing the dimensionality attribution to the cell type classification

The individual true class mean was extracted from the test cell type embedding and saved as a cache file. kNN analysis was conducted on this cache file, resulting k=5 with total 4 clusters as the most optimal clustering form. Keeping SCORE with 3,000 noising paths was run on the each true class mean – producing *l*(*c*)_*i*_, per-class log likelihood contributions of the latent space dimensionality. From this *l*(*c*)_*i*_, attributions of the each dimensionality were calculated and represented as a heatmap in Fig. 4C. The scores were computed based on the following formula (standard DDPM predictor): *s*_*θ*_(*x, t*) = − *ϵ*_*θ*_(*x, t*)*/σ*_*t*_, where 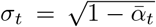. *ϵ* was not chosen based on the antithetic pairing, rather utilized completely randomly chosen values.

The permutation test was conducted to answer whether the latent dimensionality attribution patterns are more similar in within the same clusters compared to between different clusters. Pairwise Euclidean distance was calculated from the row of the heatmap as *r*_*ij*_ = ‖*row*_*i*_ − *row*_*j*_ ‖, where *row*_*i*_ = (*l*_*ci*0_, *l*_*ci*1_, *l*_*ci*2_, *l*_*ci*3_, …). For all the pairs of class i and j, two sets of distances were defined:

- *δr*_*within*_: distances (*r*_*ij*_) between the pairs of cell type classes in the same kNN-based clsuter
- *δr*_*between*_: distances (*r*_*ij*_) between the pairs of cell type classes in different kNN-based cluster

Test statistic was defined as: Δ = ⟨*δr*⟩_between_ − ⟨*δr*⟩_within_, where *<>* indicates the mean value.

The labels of the heatmap (based on KNN, k=6) was swapped for *N* = 100, 000 times. For each permutation, Δ_*perm*_ was computed to generate the null distribution. The exact p-value was calculated based on the formula: *p* − *value* = (*observations* + 1)*/*(*N* + 1) where the *observation* is |Δ_*perm*_| *>*= |Δ_*test*_ | and *N* represents the total number of permutation (two-sided test).

### Genetic perturbation classification

A diffusion model could not be effectively trained on a raw Perturb-seq dataset due to the noisy, high-dimensional, and sparse nature of the scRNA-seq dataset [53, 57, 58]. Hence, the perturbation latent space was generated by a VAE-based model with an additional perturbation-label classifier (model not provided) that was trained on a collection of Perturb-seq datasets [59–61]. This 256-dimensional latent space embedding contains subsampled 84,761 cells, spanning 4 different cell types - T cell, epithelial cell, hepatocyte, and lymphoblast. Each cell received one of the 100 unique CRISPRi-based genetic perturbations. This latent space was divided into train, validation, and test sets in 80:10:10 ratio (Table II) for the downstream task.

**TABLE II.**
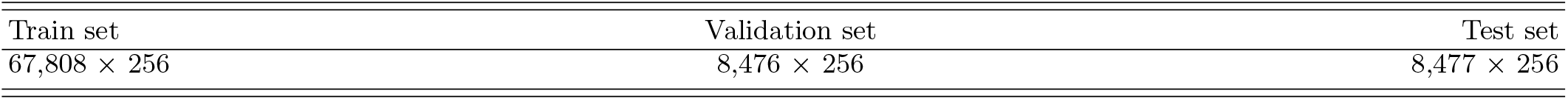
Perturb-seq embedding splits and dimensions.

| Train set | Validation set | Test set |
| --- | --- | --- |
| 67,808 × 256 | 8,476 × 256 | 8,477 × 256 |

For the attribution analysis, Keeping SCORE analysis was performed on a different 384-dimensional latent space embedding that also includes the datapoints from [62]. This embedding contains subsampled 197,249 cells with 13,992 genes, spanning 7 different cell types (cell lines) - CD4+ T cells, K562 chronic myelogenous leukemia cell line, RPE1 retinal pigment epithelia, HepG2 hepatocellular carcinoma / liver cell line, Jurkat T lymphocyte leukemia cell line, HCT116 colorectal carcinoma cell line, and HEK293T embryonic kidney cell line. Each cell received one of the 98 unique CRISPRi-based genetic perturbations (including ‘control’, without any guide RNA). This latent space was divided into train, validation, test, and out-of-distribution (OOD) datasets as described in Table III for the downstream task. The OOD dataset was not used for any analysis in this paper.

**TABLE III.** Perturb-seq attribution analysis embedding splits and dimensions.

| Train set | Validation set | Test set | OOD set |
| --- | --- | --- | --- |
| $154,240 \times 384$ | $8,570 \times 384$ | $8,567 \times 384$ | $25,872 \times 384$ |

### Genetic perturbation classification benchmark

#### 1. Keeping SCORE

Genetic perturbation mapping requires a model to effectively reconstruct the Perturb-seq transcriptional profile. To improve the quality of denoising, we designed a two-stage conditional denoiser with feature-wise linear modulation (FiLM) [63], cross-attention [64], classifier-free guidance (CFG) [65], and sigmoid beta-schedule [54] (SI Appendix, Fig. S12). The idea of a two-stage model originates from the cascaded diffusion model [66], which ramps up the image resolution by first training a smaller model and then passing its output to a larger model. Similarly, we designed a pseudo two-stage architecture within a single training loop, where the low-dimensional MLP (512 dimensions) first captures a coarse representation of the data, and then learns fine-grained details at the higher dimensionality stage (2048 dimensions). In stage 1, the model learns coarse and global structure using 512-dimensional MLP layer with layer normalization and feature-wise cross-attention. The second stage expands to 2048 dimensions, applying the similar sequence but replacing cross-attention with FiLM to produce the final refined prediction. The final output is a residual sum of the two predictions. We encoded 1,000 timesteps (encoded by sinusoidal embedding [56]) and perturbation labels into conditional embedding and concatenated into a single condition vector that is passed at the start of each stage. Beta scheduling was also designed to maximize the conditioning effect by employing the sigmoid schedule from the scVAEDer model [53] to learn the fine-grained details [54]. The maximum beta value was capped to 0.999 for numerical stability. The CFG probability of 0.2 was applied to enhance the conditioning effect.

The diffusion model was trained and validated on the Perturb-seq embedding. The training loop was set to 500 epochs with early-stopping based on the validation loss with an improvement threshold of 10^−5^, and patience of 30. For our model, the training was early-stopped after 113 epochs of training. The learning rate was scheduled based on the cosine decay with the warm-up rate of 5%. The denoised UMAP was generated for every 30 epochs based on the EMA weights. After training, the fully-trained diffusion model was passed to Keeping SCORE with 4 noising paths. Despite the model being trained based on both raw weights and exponential moving average (EMA)-based weights, only the raw weights were used for the Keeping SCORE classification task. The model configuration and training details can be found in Table S4. The loops in the Keeping SCORE were reshaped, vectorized, and run on GPU to resolve the computational bottleneck. The output from the diffusion model network was directly used as the score function value. Keeping SCORE’s structure utilized anthitetic pairing, meaning that for every noise *ϵ*, the paired − *ϵ* was included to improve Monte Carlo estimation stability. For the uncertainty aware analysis, we considered model’s prediction was correct when the highest classified class error bar (the standard error of each noise path) contains the true class mean.

#### 2. Multi-layer Perceptron

A simple MLP was built with 3 hidden layers with 512, 256, 128 dimensions each. The MLP model was trained on the full train embedding loaded without any additional subsampling or modification. The validation loss was calculated based on the validation embedding. For MLP, the training was early-stopped after running 25 epochs out of 100. The model’s performance was then evaluated on the full test embedding. The model structure and training details can be found in Table S5.

#### 3. XGBoost

CPU-based XGBoost (version 3.1.1) was used for genetic perturbation mapping. The XGBoost model was trained on the full train embedding without any additional subsampling or modification. The validation loss was calculated based on the validation embedding. For XGBoost, training was stopped after running the full 500 epochs. The model’s performance was then evaluated on the full test embedding. The non-default model hyperparameters and training details can be found in Table S6.

#### 4. Logistic Regression

We used a logistic regression model from Scikit-learn (version 1.7.2) for genetic perturbation mapping. The logistic regression model was trained on the full train embedding without any additional subsampling or modification. The validation embedding was not utilized as the classic logistic regression model does not support the early-stop. The model’s performance was then evaluated on the full test embedding. The non-default model hyperparameters and training details can be found in Table S7.

### Computing the attributions in perturbation classification

The embedding used for the Perturb-seq attribution analysis contains additional data from Zhu et al. (2025) [62]. Accordingly, a diffusion model was trained on the 384 input embedding dimensions (SI Appendix, Fig. S9 for the embedding and SI Appendix, Fig. S13 for the model structure). In the first stage, two 512 dimensional hidden layers are used with layer normalization, attention layer, FiLM layer, and GELU activation function. The dropout of 0.1 was included between the two hidden layers, after the first GELU activation function. The second stage resembled the first layer with larger dimensions (two 2,048 dimensional hidden layers + layer normalization + attention layer + FiLM layer + GELU activation function with dropout of 0.1 after the first activation function). The parameters of the model were identical to the parameters described in classification task (SI Appendix Fig. S4, except for the latent dimensionality = 384 and noise path = 10,000). The diffusion model was trained with the complete training set and validated with the validation set. The training run early-stopped at 145 epochs after 30 epochs of patience, reporting the final training loss of 0.090097 and validation loss of 0.065373. The individual true class mean was extracted from the Perturb-seq test embedding and saved as a cache file. kNN analysis was conducted on this cache file, resulting k=6 as the most optimal clustering form. Keeping SCORE with 10,000 noising paths was run on the each true class mean – producing *l*(*c*)_*i*_, per-class log likelihood contributions of the latent space dimensionality. Then these *l*(*c*)_*i*_ was decoded using the linear decoder. From these decoded *l*(*c*)_*i*_, the attributions for each dimensionality were calculated and represented as a heatmap in Fig. 4D and individual values in Fig. 4E. The scores were computed based on the following formula (standard DDPM predictor): *s*_*θ*_(*x, t*) = −*ϵ*_*θ*_(*x, t*)*/σ*_*t*_, where 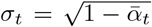 [56]. Keeping SCORE *ϵ* was not chosen based on antithetic pairing, rather utilized completely randomly chosen values. We applied a learned linear decoder to compute Δ*g*_*i*_ the perturbation responses from the latent space to the gene space. Initially, in the latent space, we produce 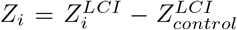, where i indicates the individual perturbation types, control representing the control cells, and LCI as the incremental log-likelihood. The linear decoder (*W*_*dec*_ = *W*_2_*W*_1_, where *W*_1_ is the fully-connected layer weights and *W*_2_ is the gene loading matrix of the decoder) can then map the gene to the gene space as following:

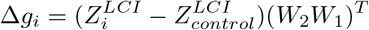

Practically, for the latent space k and the perturbation i and gene j, 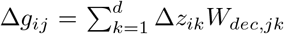, where *W*_*dec,jk*_ = (*W*_2_*W*_1_)_*jk*_.

### Predicting how mutations change protein stability

We start with FireProtDB [40] entries with measured ΔΔ*G* and associated metadata, filtering to rows with ΔΔ*G*, full amino-acid sequence, wild-type and mutant residues, mutation position, pH, ASA, B-factor and conservation score. For each mutation, we extract a local window of length 11 amino acids centered on the mutation site (5 residues on each side, truncated at sequence ends if needed). We tokenize using a 21-symbol vocabulary (20 amino acids and a padding index 0), concatenate WT and mutant windows into a single token sequence of length 22. We then normalize the four covariates to a range of 0 to 1 using MinMax scaling.

The conditional autoencoder is comprised of an encoder and linear decoder. Prior to an MLP encoding into a 3-dimensional latent vector, the tokens are embedded into a learned 32 dimensional embedding space, passed through a 2-layer transformer with multi-head self attention and mean-pooled to obtain a single sequence embedding. In the latent space, we have a direct regression head where the latent variable *z* is passed through a deep residual MLP that outputs a scalar prediction for ΔΔ*G*.

In the latent space, we have a forward, VP, latent diffusion process over *z* with a linear noise ramp between 10^−4^ and 0.002. Time and ΔΔ*G* are embedded with separate MLPs and concatenated with *z* before being passed through a linear layer plus multiple residual blocks and a final linear layer outputting the predicted noise at that time and position.

We perform gradient descent directly on the likelihood *l*(ΔΔ*G*) using automatic differentiation, noising each test point to calculate it over the *T* = 2 diffusion schedule to calculate *l*, and taking *N* = 20 small steps weighted by the learning rate *α* = 0.1. We clamp the updated estimates of ΔΔ*G* at [− 15, 20] to stabilize the gradient descent. From here, since every step of the gradient descent is itself differentiable we use the mean-squared error between the true and estimated ΔΔ*G* as our loss to train the network to infer ΔΔ*G* in the chosen number of steps, and weight this loss strongly during training.

Our total loss function is a combination of reconstruction loss, classical mean-squared error regression loss, and the diffusion classification loss, weighted with constants *λ*_*recon*_ = 1.0, *λ*_*reg*_ = 0.25, *λ*_*Diff*_ = 10.0.

We train the network using AdamW with a learning rate of 10^−4^, and batch size of 256 for 2500 epochs.

For the mega-scale protein stability dataset [41], we use the same model architecture, and train/evaluate it on two separate train, test splits. The first splits the data by mutation whereas the ladder splits the data by protein[42], so if more conservative. The experimental covariates include predicted protease-resistance metrics under trypsin and chymotrypsin digestion, a model-inferred absolute folding free energy, and an experimentally inferred absolute folding free energy.

## ACKNOWLEDGEMENTS

We thank members of the Goyal lab, especially Madeline E. Melzer, for helpful discussions and comments on the manuscript. We thank the Feinberg Information Technology, Northwestern University Information Technology, and Quest High Performance Computing Cluster at Northwestern University Feinberg School of Medicine for their services. We acknowledge support from the National Institute for Theory and Mathematics in Biology (NITMB) through the National Science Foundation (DMS-2235451) and the Simons Foundation (MPTMPS-00005320). BKS acknowledges support from NUCATS T32, NITMB. JJ, CP and LS were supported by funding from YG. S.V. was supported by the National Institute of General Medical Sciences of the NIH under Award No. R35GM147400. YG acknowledges support from the Pew-Stewart Scholars Program, Burroughs Wellcome Fund Career Awards at the Scientific Interface, as well as AHA (24CSA1256987), and Edward Mallinckrodt Jr. Foundation. YG is a CZ Biohub Investigator. This research benefited from the National Science Foundation through the Center for Living Systems (grant no. 2317138) We acknowledge our protein cartoons in Figures 1 and 5 were created in BioRender. Kuznets-Speck, B. (2025) https://BioRender.com/fbbwsvn (Licensed under CC BY 4.0).

## AUTHOR CONTRIBUTION

BKS and YG conceived and designed the study. BKS designed and performed analyses with input from YG. JJ performed computations for the genomics examples with BKS. LS collected and harmonized the datasets, and prepared the code repository. PP assisted with figures and manuscript preparation. AZ and SV assisted in formulating the theoretical framework, and EP on machine learning. BKS and YG wrote the manuscript with input from all authors.

## AUTHOR DECLARATION

YG receives consultancy from Schmidt Sciences and Gilead Sciences.

## DATA AND CODE AVAILABILITY

This paper analyzes primarily pre-existing, publicly available datasets. The datasets can be accessed through google drive (see github README for link).All code for the analyses in this manuscript has been deposited at: https://github.com/GoyalLab/KeepingScore.

## MNIST

We trained our convolutional VAE–classifier model on the MNIST handwritten digit dataset (LeCun et al., 1998) [67], downloaded automatically via torchvision.datasets.MNIST.

## CIFAR10

Embeddings were generated using the penultimate activations of a ResNet-18 pretrained on ImageNet-1k (torchvision model zoo), applied to CIFAR-10 images available at https://www.cs.toronto.edu/∼kriz/cifar.html [24].

### Genomics

For the cell type classification task, we utilized and benchmarked codes from Fischer et al. (2024) [25] available at https://github.com/theislab/scTab/. Perturb-seq dataset for the benchmark analysis was collected from Peidli et al. (2024) [59] available at: https://projects.sanderlab.org/scperturb/. From there, we downloaded Replogle et al. (2022) [33] and Nadig et al. (2025) [61]. In addition, the original Huang et al. (2025) [60] dataset was processed and included. For the Perturb-seq attribution anlaysis, large scale CD4+ T cells from Zhu et al. (2025) [62] was added.

### Protein

We used experimentally curated protein stability measurements from FireProtDB https://loschmidt.chemi.muni.cz/fireprotdb/ [40], a database of ΔΔG values for point mutations in proteins, and processed the sequences and metadata directly from the FireProtDB CSV export.

## Supplementary Information

## Appendix 1. Runtime analysis for different likelihood estimators

We illustrate the relative computational efficiency of the backwards Itô scheme in SI Appendix Fig. S1, where we show the runtime per trajectory for a *T* = 1, 000 diffusion process under the forward and backwards Itô estimators along with that from a probability flow ODE estimator for gaussian mixture models of increasing dimension. The forward Itô and probability flow ODE estimators are dominated by the score-divergence term for large dimensions.

**FIG. S1.**
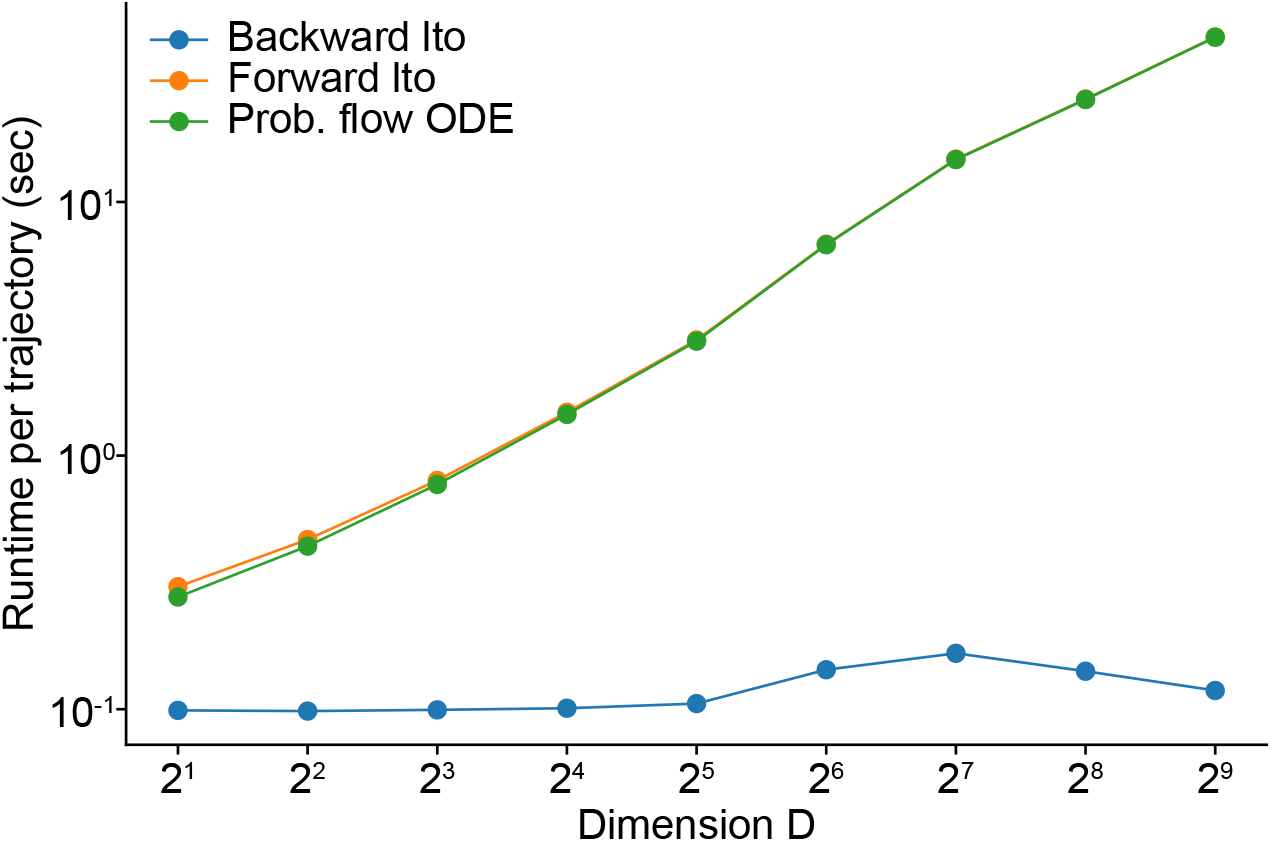
Runtime scaling for diffusion likelihood estimators with (forward Itô, prob. flow ODE) and without (backward Itô) the score-divergence term.

## Appendix 2 Details about the Keeping SCORE time step subsampling analysis

Subsampling the timestep from a complex non-linear space is unintuitive. Both of our diffusion models for genetic analysis are based on sigmoid timestep scheduling. To successfully shrink the timestep while maintaining its original features, we subsampled based on the *β*-space itself instead of the timestep-space. This idea is similar to the timestep subsampling concepts in Denoisning diffusion implicit model (DDIM) [68]. Specifically, we compute all the cumulative beta values across the full schedule, then sample the new timesteps uniformly corresponding to the cumulative beta level. Each *β*-level is mapped back to its closest timestep, maintaining the overall sigmoid cumulative *β* feature, preserving the structure of the *β* trajectory.

**FIG. S2.**
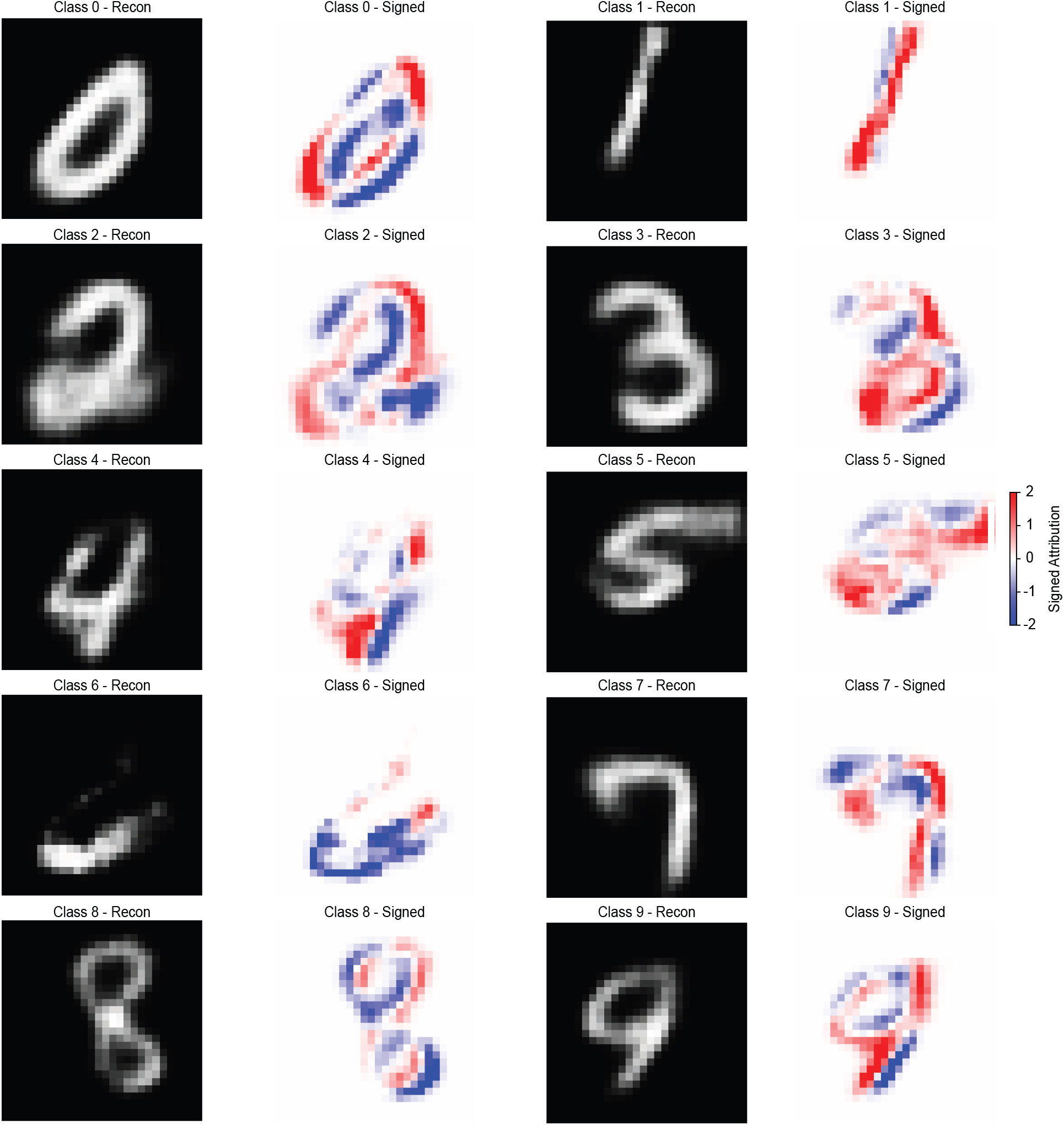
Testing reconstructions from VAE latent space and accompanying signed attribution to classification, one randomly selected for each digit

**FIG. S3.**
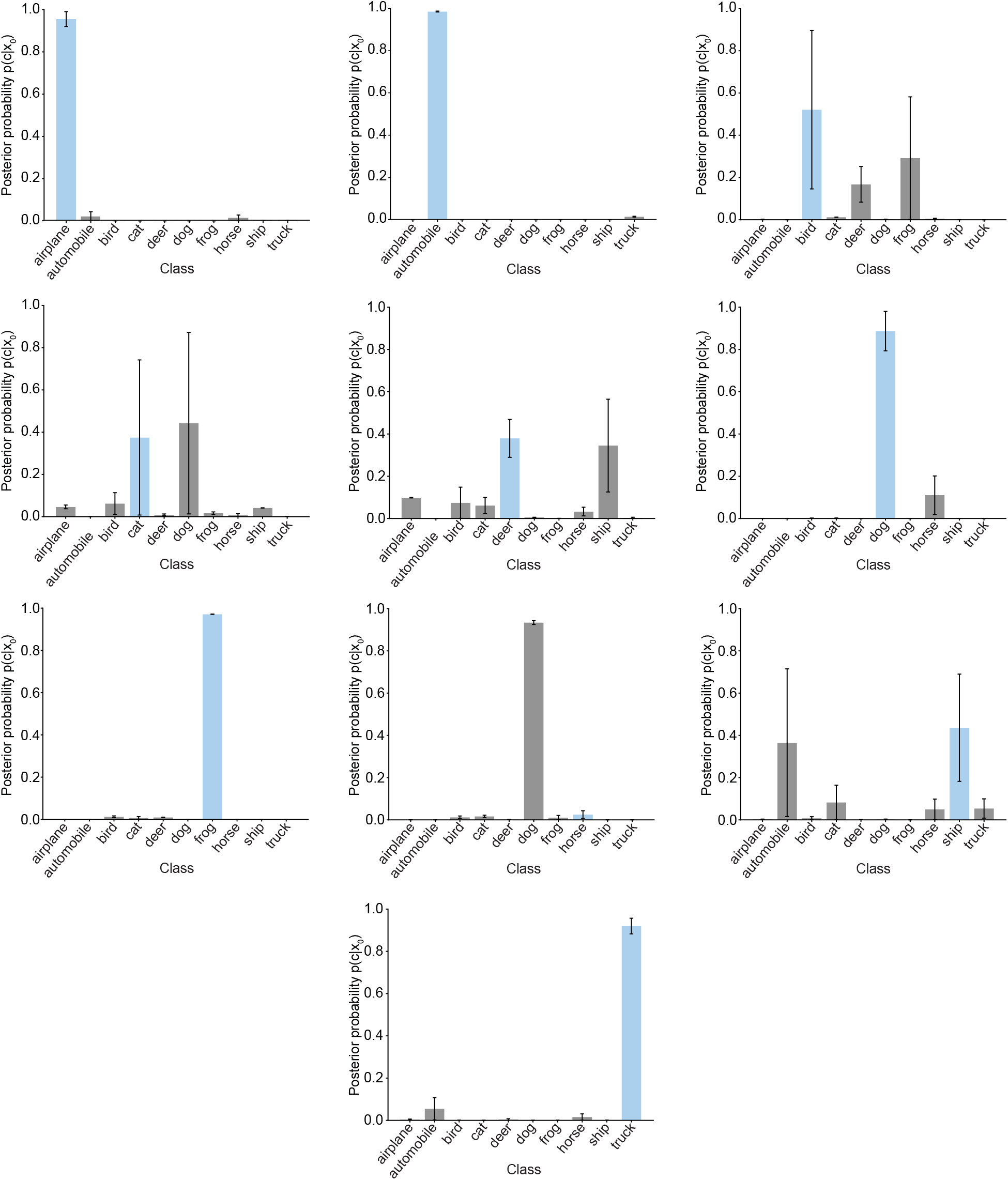
Posterior probabilities for CIFAR test embeddings, one from each class. The sky blue color columns represent the correct class and the black vertical on top of each column represents the error bar of each estimation. Keeping SCORE correctly mapped most of the images.

**FIG. S4.**
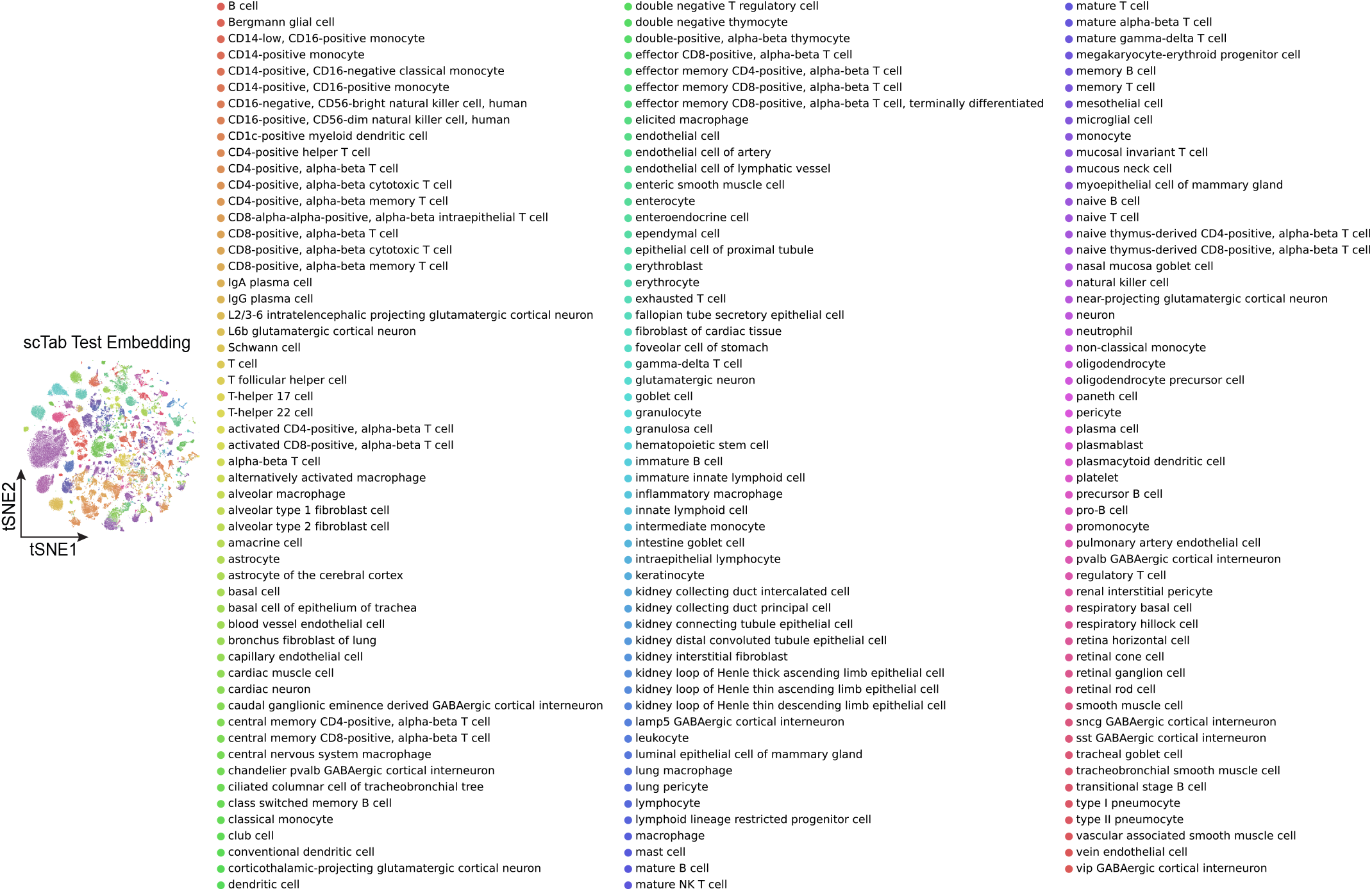
tSNE plot of the cell type test embedding with labels. The embedding is obatined from scTab data and total of 200,000 cells are subsampled for the illustration purpose. Overall, immune cells dominate with multiple T cell and B cell subtypes.

**FIG. S5.**
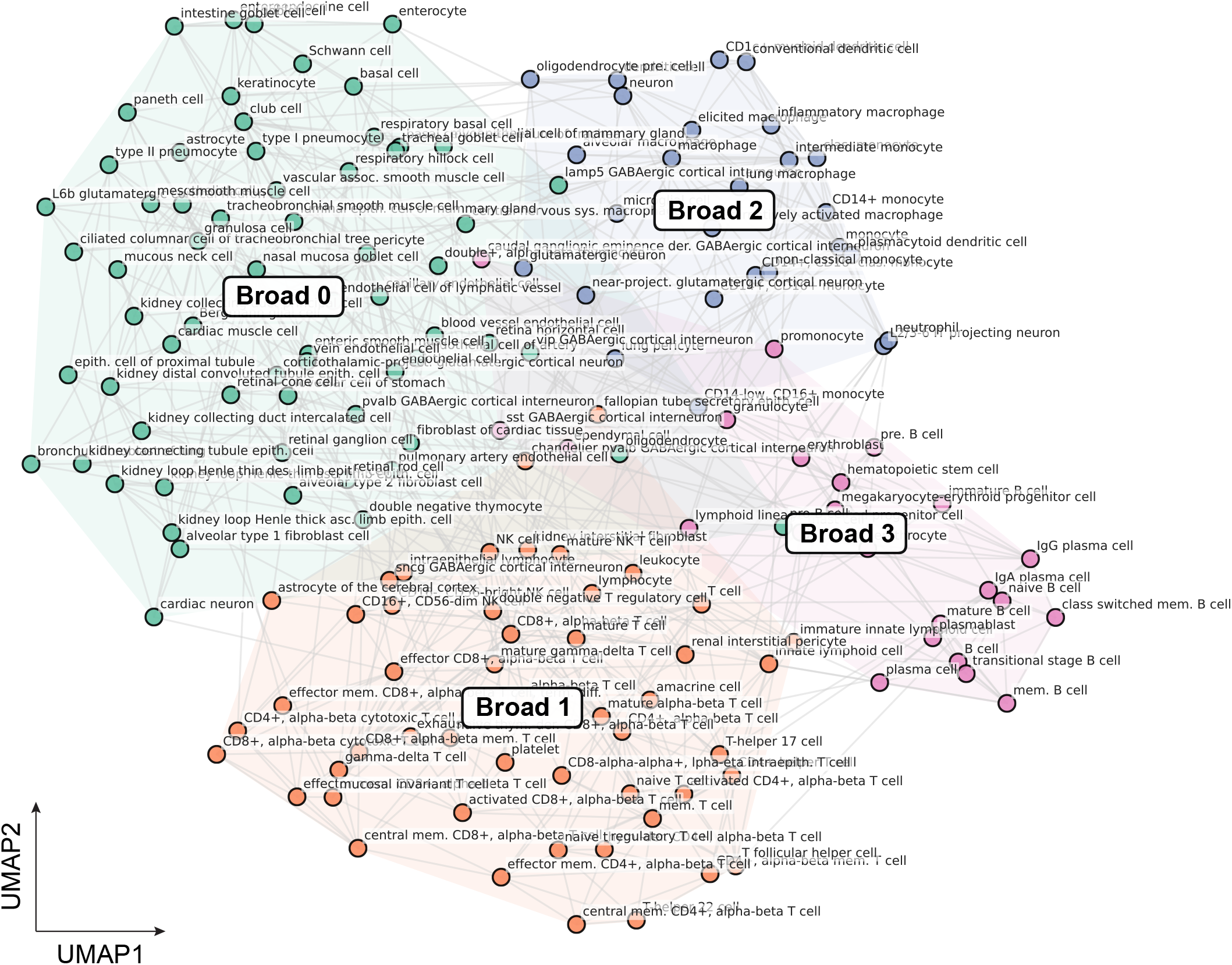
UMAP plot of kNN clustering on the mean of each cell type class. K=5 with the 4 broad cell type clusters was chosen. This map is the basis for the clusters in Fig. 3C. Similar cell types are clustered together - for example, T cells tend to sit in Broad1 cluster whereas B cells are assigned to Broad2 cluster.

**FIG. S6.**
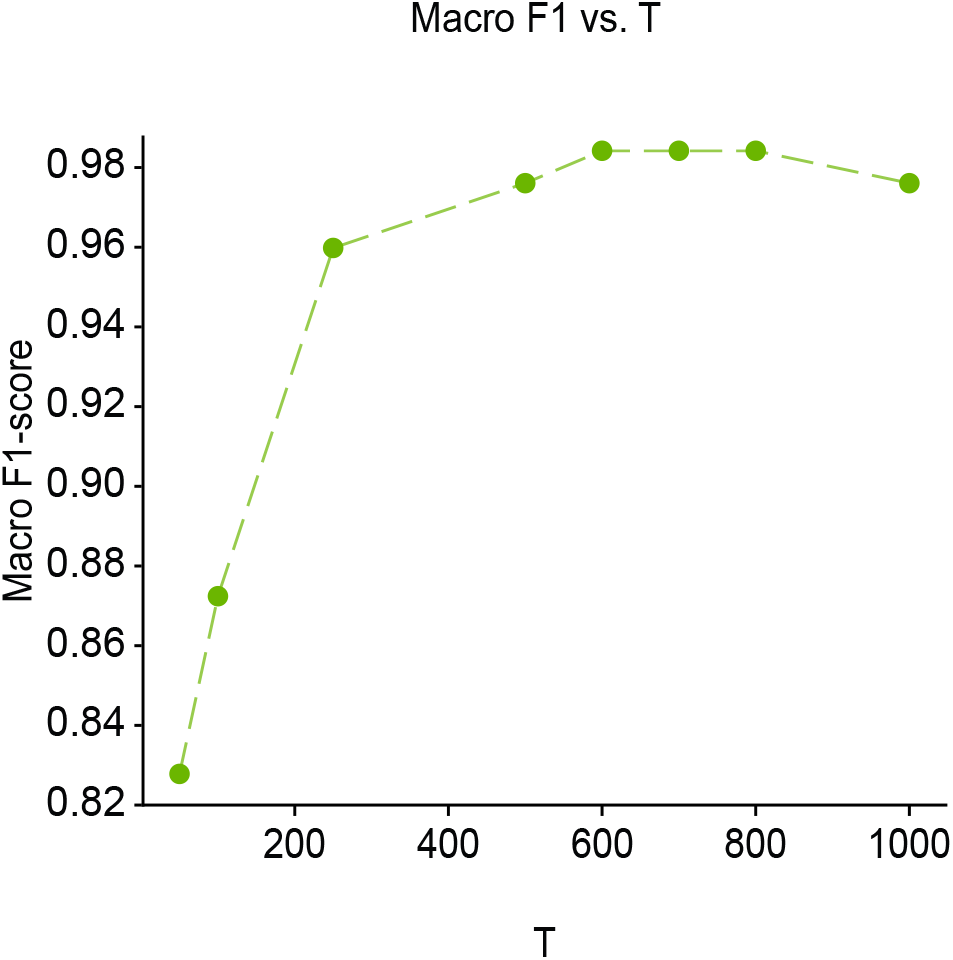
The plot shows the diffusion model’s macro-average F1-score changes along the subsampled Keeping SCORE inference timestep (see Material Methods and SI Appendix 1; the model was trained with 1,000 steps). The macro-average F1-score is calculated based on the mean gene expression level of each cell type class (1 support for each class). Around the timestep of 600, the model’s macro-average F1-score gets almost saturated, and the original macro-average F1-score reaches 0.984 (not uncertainty aware). This shows that running with the fewer Keeping SCORE inference timesteps can reduce the computational time while achieving the comparable analysis result.

**FIG. S7.**
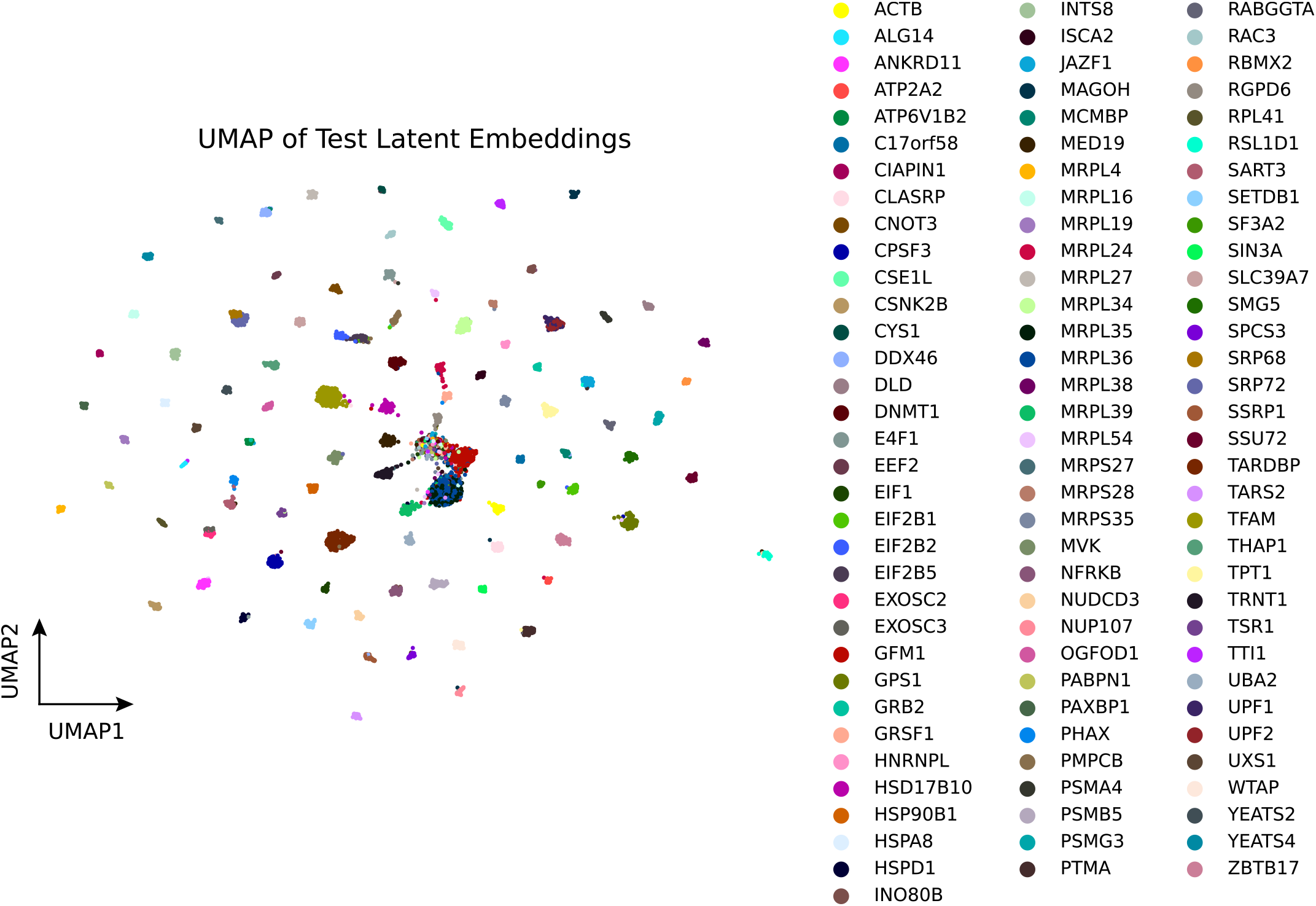
The UMAP visualization of the Perturb-seq test embedding with labels. Each colors represent the 100 different perturbation types. All the cells in the test embedding was used for the plot. Although most of the perturbation types are well separated across the latent manifold, the center of the embedding is blurred due to the similarity between perturbations.

**FIG. S8.**
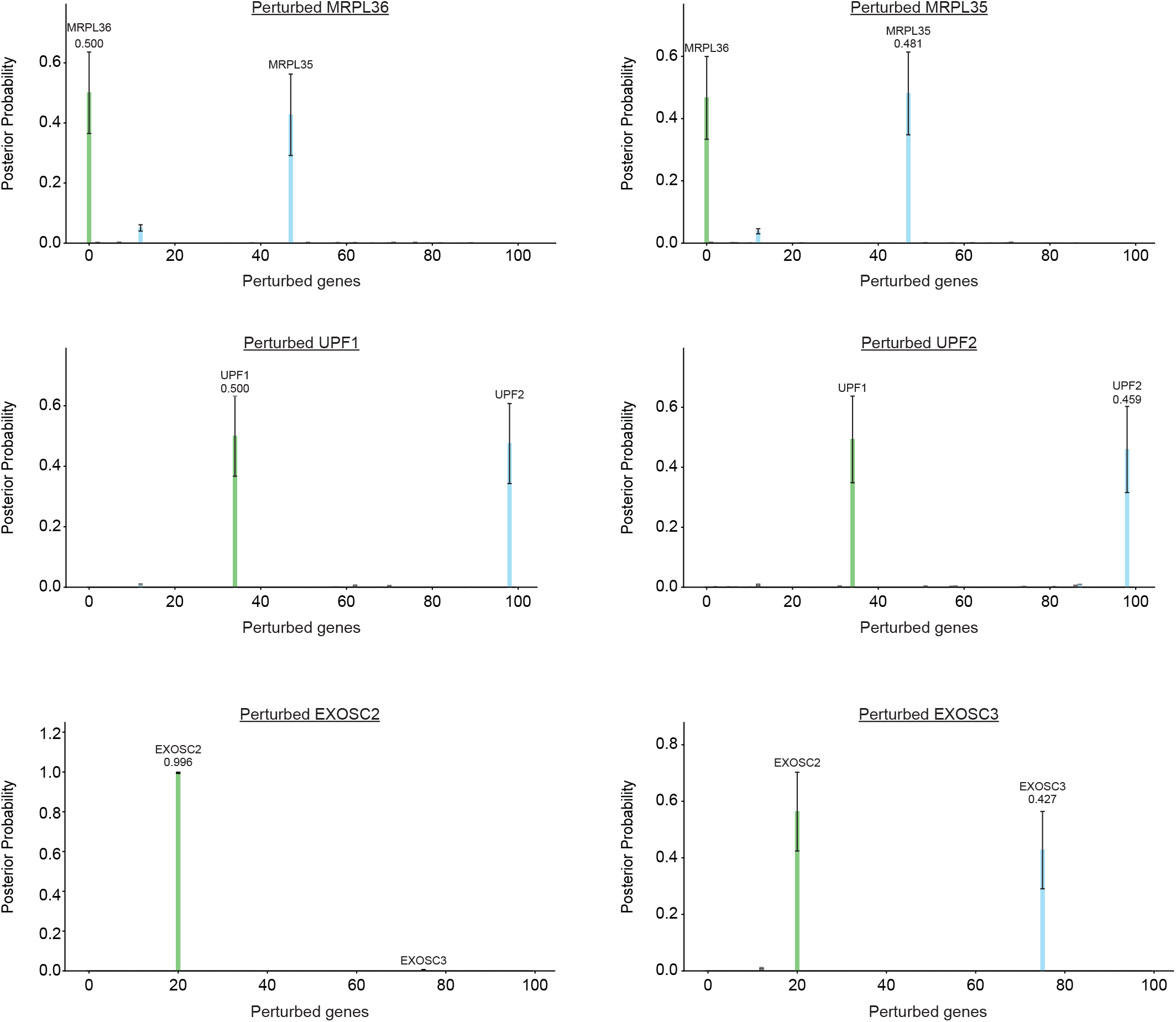
Additional tie match cases where the biological relationships are symmetrical and asymmetrical. (A) Symmetric tie match case observed in MRPL36 and MRPL35. Although perturbation of MRPL36 is correctly assigned to MRPL36 (0.500 ± 0.136), it is also frequently mapped to MRPL35 (0.427 ± 0.135) with the overlap of error bars. The similar case was observed when classifying MRLP35 perturbation (MRPL35: 0.481 ± 0.133 followed by MRPL36: 0.466 ± 0.133). MRPL35 and MRPL36 are both components of the mitochondrialribosome subunit 39S [69]. (B) Additional symmetric tie match observed in UPF1 and UPF2. UPF1 and UPF2 forms heterodimer for the activation of nonsense mediated decay (NMD) [70]. UPF1 was mapped to UPF1 (0.500 ± 0.132) with UPF2 as the second best prediction (0.475 ± 0.132). UPF2 was assigned to UPF1 (0.493 ± 0.133) followed by mapped to UPF2 (0.459 ± 0.144). (C) Tie match cases where the biological relationships are asymmetrical. EXOSC2 is mostly mapped to EXOSC2 (0.919 ± 0.015), but EXOSC3 is frequently mapped to EXOSC2 (0.396 ± 0.126). EXOSC2 and EXOSC3 proteins are both involved in forming nuclear RNA exosome trimeric cap [71].

**FIG. S9.**
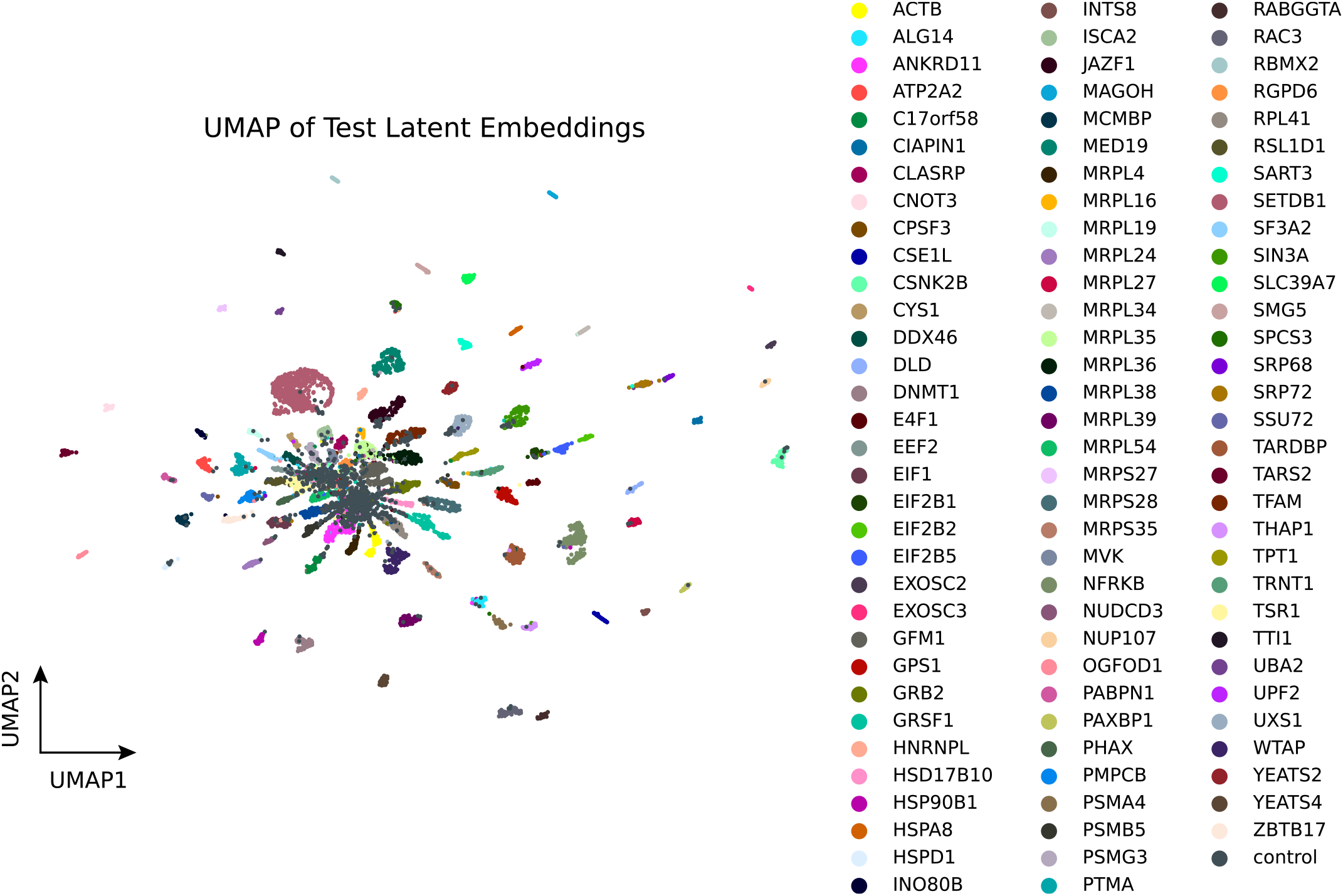
UMAP visualization of the test embedding of Perturb-seq data used for the attribution analysis. All the cells in the test embedding was used for the plot. Each colors represent 98 perturbation types (including ‘control’). Although most of the perturbation types are well separated across the latent manifold, the center of the embedding is blurred by the control cells.

**FIG. S10.**
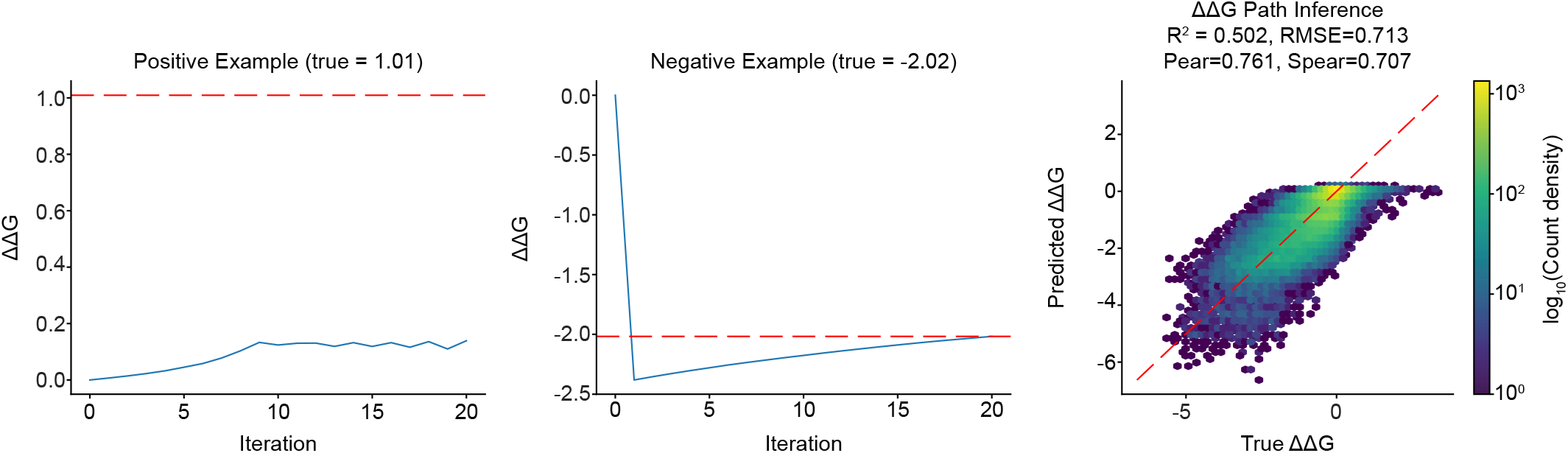
Keeping SCORE predicts protein stability changes from mega-scale measurements of protein stability, split by mutation. (Left/Center) Inferred ΔΔ*G* as a function of gradient descent iteration for a stabilizing and destabilizing mutation. (Right) True and Inferred ΔΔ*G* histogram for the entire test set.

**FIG. S11.**
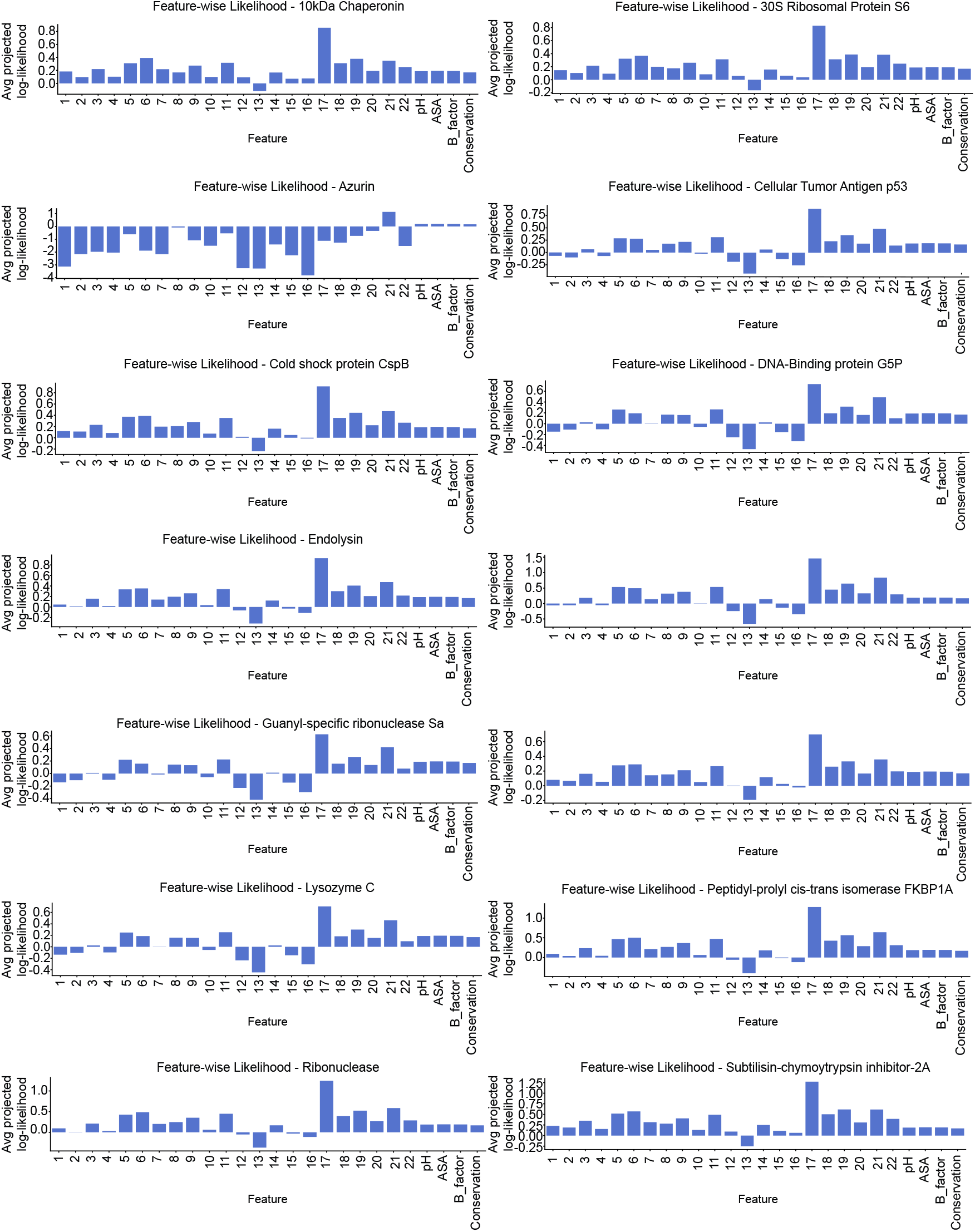
Posterior feature-wise log-likelihood for classification averaged over mutations for specific proteins.

**FIG. S12.**
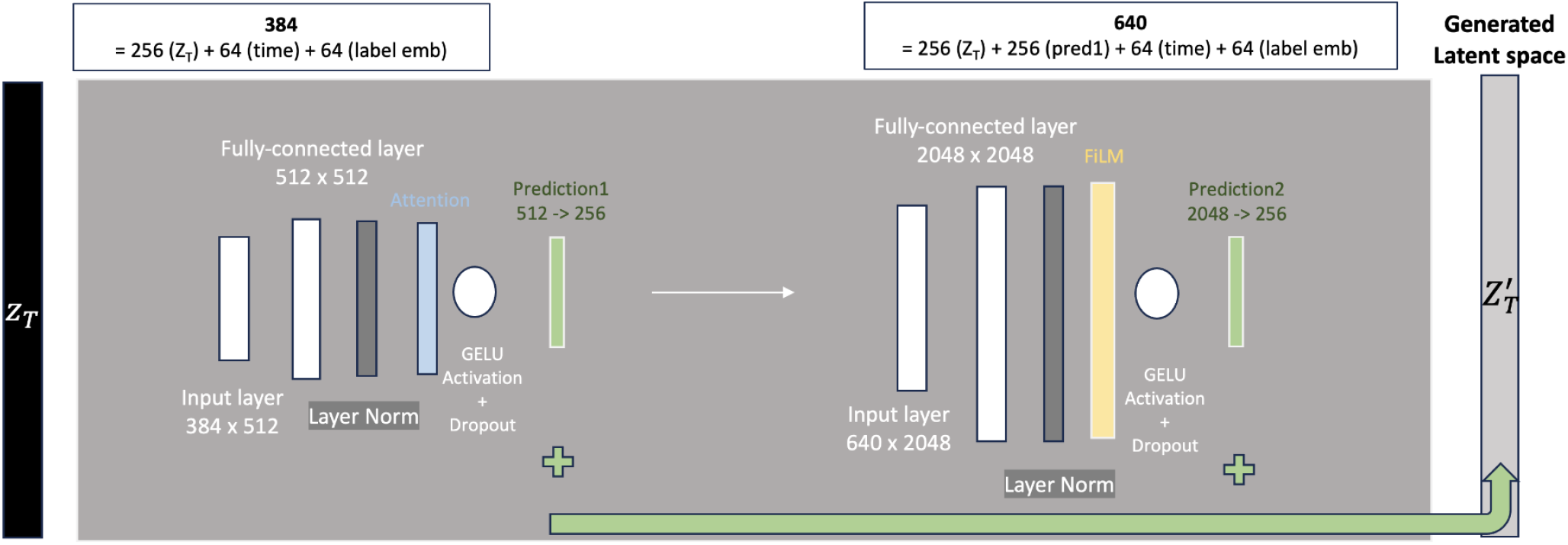
The schematic description of Perturb-seq diffusion model architecture. The Keeping SCORE result in Fig. 4B was generated based on this diffusion model. To capture the subtle signals from CRISPRi-based genetic knockdown, our diffusion model employed a complex structure to provide powerful conditioning effect. *Z*_*T*_ indicates a completely noised input sample (complete Gaussian noise) after the forward process (noising steps) of the diffusion model. The model is cascaded into the two similar steps - (I) the first block with the smaller 512 dimensional hidden layer, (II) and the second block with the larger 2,048 dimensional hidden layer. The input layer contains the input (*Z*_*T*_), timestep dimension (64 dimensions used), and label embedding dimension (64 dimensions; conditioning), adding up to 384 in total (upper left white box). This input is passed to hidden layer (white bar) → fully connected 512 dimensional hidden layer (white bar) → layer normalization (dark grey bar) → attention layer (sky blue bar) → GELU activation function with dropout rate of 0.1 (white circle). The first prediction is 256 dimensions (green column). The second predictions follow the similar steps, with the 2,048-dimensional hidden layer. Instead of the attention layer in the first prediction, second prediction incorporated the FiLM layer (yellow bar). The second prediction was also transformed to 256 dimensions (green bar). Two predictions are concatenated as prediction1 + prediction2 (green arrow) to generate the final reconstructed latent space 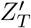. This reconstructed latent space is not decoded.

**FIG. S13.**
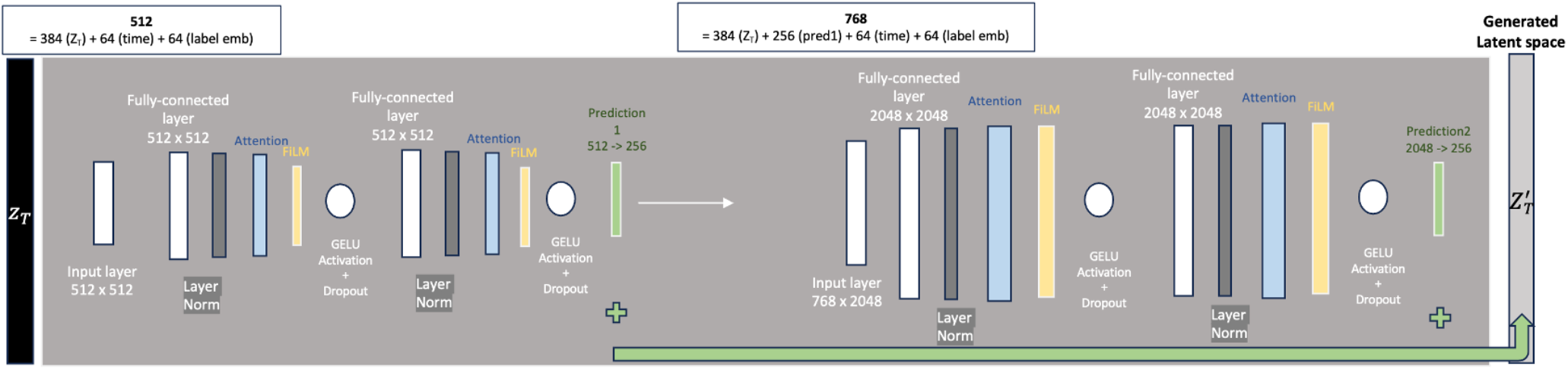
The schematic description of Perturb-seq diffusion model used for the attribution analysis. The Keeping SCORE result in Fig. 4D and E were generated based on this diffusion model. The overall structure is similar to SI Appendix, Fig. S11. However, two hidden layers are used in each step instead of one, and FiLM layers and attention layers are applied after each hidden layer. The latent space was decoded by a linear decoder described in Material and Methods.

**TABLE S1.** Cell type classification: KeepingSCORE hyperparameters.

| Parameter | Value |
| --- | --- |
| **Sigmoid Beta Schedule Parameters** |  |
| T | 1000 |
| beta_min | $10^{-5}$ |
| beta_max | 0.022 |
| low | -6 |
| high | 6 |
| **Model Configuration** |  |
| latent_dim | 128 |
| time_dim | 64 |
| label_emb_dim | 64 |
| hidden_dim | 512 |
| **Training Configuration** |  |
| batch_size | 2048 |
| epochs | 500 |
| lr | $3 \times 10^{-4}$ |
| lr_finder min_lr | $10^{-6}$ |
| lr_finder max_lr | $10^{-2}$ |
| lr_finder num_training | 100 |
| optimizer | AdamW |
| adam_betas | (0.9, 0.9999) |
| wd_model | $10^{-7}$ |
| patience | 25 |
| **KeepingSCORE Setting** |  |
| noise paths | 4 |

**TABLE S2.** Cell type classification: XGBoost hyperparameters.

| Parameter | Value |
| --- | --- |
| **Model Configuration** |  |
| n_estimators | 1000 |
| eta | 0.075 |
| max_depth | 10 |
| subsample | 0.75 |
| objective | multi:softprob |
| eval_metric | mlogloss |
| tree_method | gpu_hist |
| **Training Specification** |  |
| early_stopping_rounds | 10 |
| n_jobs | 20 |
| sample_weight | weights |
| eval_set | [(x_val, y_val)] |

**TABLE S3.** Cell type classification: Logistic Regression hyperparameters.

| Parameter | Value |
| --- | --- |
| loss | log |
| random_state | 1 |

**TABLE S4.**
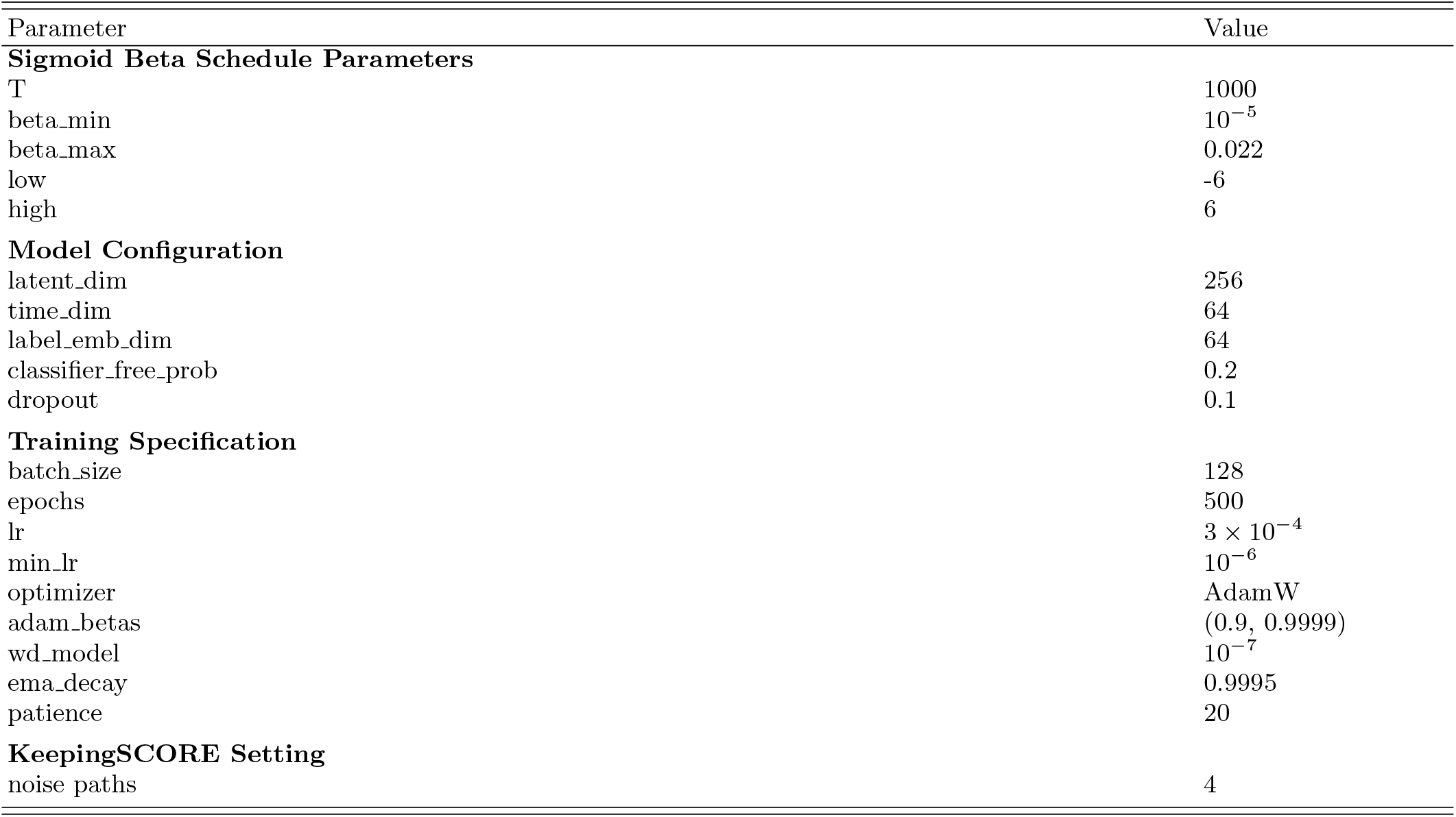
Perturb-seq: KeepingSCORE hyperparameters.

**TABLE S5.** Perturb-seq: Multi-layer Perceptron hyperparameters.

| Parameter | Value |
| --- | --- |
| **Model Structure** |  |
| hidden_dims | [512, 256, 128] |
| dropout | 0.3 |
| **Training Specification** |  |
| batch_size | 128 |
| epochs | 100 |
| lr | $10^{-3}$ |
| lr_scheduler | ReduceLROnPlateau |
| lr_scheduler:factor | 0.5 |
| lr_scheduler:patience | 5 |
| optimizer | Adam |
| weight_decay | $10^{-4}$ |

**TABLE S6.** Perturb-seq: XGBoost hyperparameters.

| Parameter | Value |
| --- | --- |
| **Model Non-default Hyperparameters** |  |
| eta | 0.1 |
| objective | multi:softprob |
| num_class | 100 |
| eval_metric | mlogloss |
| tree_method | hist |
| **Training Specification** |  |
| num_boost_round | 500 |
| early_stopping_rounds | 20 |
| evals | [(dval, "eval"), (dtrain, "train")] |

**TABLE S7.** Perturb-seq: Logistic Regression hyperparameters.

| Parameter | Value |
| --- | --- |
| random_state | 42 |
| max_iter | 100 |
| multinomial | auto |

